# Recovery of human upper airway epithelium after smoking cessation is driven by a slow-cycling stem cell population and immune surveillance

**DOI:** 10.1101/2025.08.26.672394

**Authors:** Hugh Selway-Clarke, Kate H. C. Gowers, Vitor H. Teixeira, Calum Gabbutt, Carlos Martínez-Ruiz, Ahmed S. N. Alhendi, Benjamin D. Simons, Benjamin A. Hall, Nicholas McGranahan, Anob M. Chakrabarti, Sam M. Janes, Adam Pennycuick

**Affiliations:** Lungs for Living Research Centre, UCL Respiratory, University College London, London, UK; I-X Centre for AI in Science, Imperial College London, London, UK; Centre for Evolution and Cancer, Institute of Cancer Research, London, UK; UCL Cancer Institute, University College London, London, UK; Cancer Evolution and Genome Instability Laboratory, The Francis Crick Institute, London, UK; The Wellcome Trust/Cancer Research UK Gurdon Institute, University of Cambridge, Cambridge, UK; Department of Medical Physics and Biomedical Engineering, University College London, London, UK; University College London Hospitals NHS Trust, London, UK

## Abstract

The upper airway epithelium in humans is maintained in homeostasis by a resident population of basal stem cells. In the presence of tobacco smoke these gain mutations that significantly increase their risk of transformation to lung squamous cell carcinoma. Previous studies show that a small proportion of stem cells avoid the mutational damage caused by carcinogens in tobacco and are more abundant in the lungs of former smokers than ongoing smokers, indicating unexplained tissue-level genomic recovery. This mirrors epidemiological risk, which falls rapidly after quitting smoking. Somatic evolutionary mechanistic hypotheses have been proposed to explain these observations. Here, we present a computational framework to model each of these hypotheses within the upper airway epithelial stem cell population over the entire patient lifetimes of a cohort with diverse smoking histories. Applying a mechanistic learning approach based on a set of biologically informed metrics to single cell-derived whole-genome sequencing data, we identified subtle differences between epithelia modelled under different combinations of hypotheses. A slow-cycling subpopulation of stem cells, combined with suppression of immune predation of highly mutated stem cells while smoking, best matched observed data, a result converged upon by multiple distinct machine learning methodologies. Our findings, drawing on an evolutionary model of mutagen exposure at a whole-lifetime scale that is not feasible to model in vivo, reveal the mechanisms behind reduction in lung squamous cell carcinoma risk on cessation of smoking and inform future therapeutic interventions to prevent lung cancer initiation.

## INTRODUCTION

Smoking causally increases lung cancer risk^1^ and tobacco smoke is a known mutagen^2^. Healthy somatic tissues in the human body are patchworks of clones^3–11^, subject to evolutionary processes that act on mutations arising from exogenous and endogenous mutational processes. The accumulation of somatic mutations is a leading candidate cause for many cancers, due to the significant differences in the genomes of healthy and cancerous tissue^6,12^. In addition to being mutagenic^2^ and activating^13^, carcinogens can also theoretically work as selectogens: altering the fitness landscape to benefit mutated cells without directly driving them to cancerous transformation^14,15^.

Two recent whole genomic sequencing (WGS) studies using cell cultures derived from individual basal cells taken from samples of healthy human upper airway epithelial tissue^3,4^ from cohorts with diverse smoking histories (Figure 1A) have delineated a gradual accumulation of mutational burden over the course of life, accelerated by tobacco smoking (Figure 1B). However, within-patient heterogeneity of mutational burden among patients with a history of smoking revealed two surprising dynamics: first, the mutational burden of airway basal cells had an approximately bimodal distribution in many patients with a history of smoking, with some cells near normal while others were highly mutated (Figures 1C, 1D). Moreover, the proportion of near-normally mutated cells was higher in ex-smokers than in current smokers^3^ (Figures 1C, 1D), suggesting that this subpopulation of cells expands proportionally after cessation of smoking.

**Figure 1.**
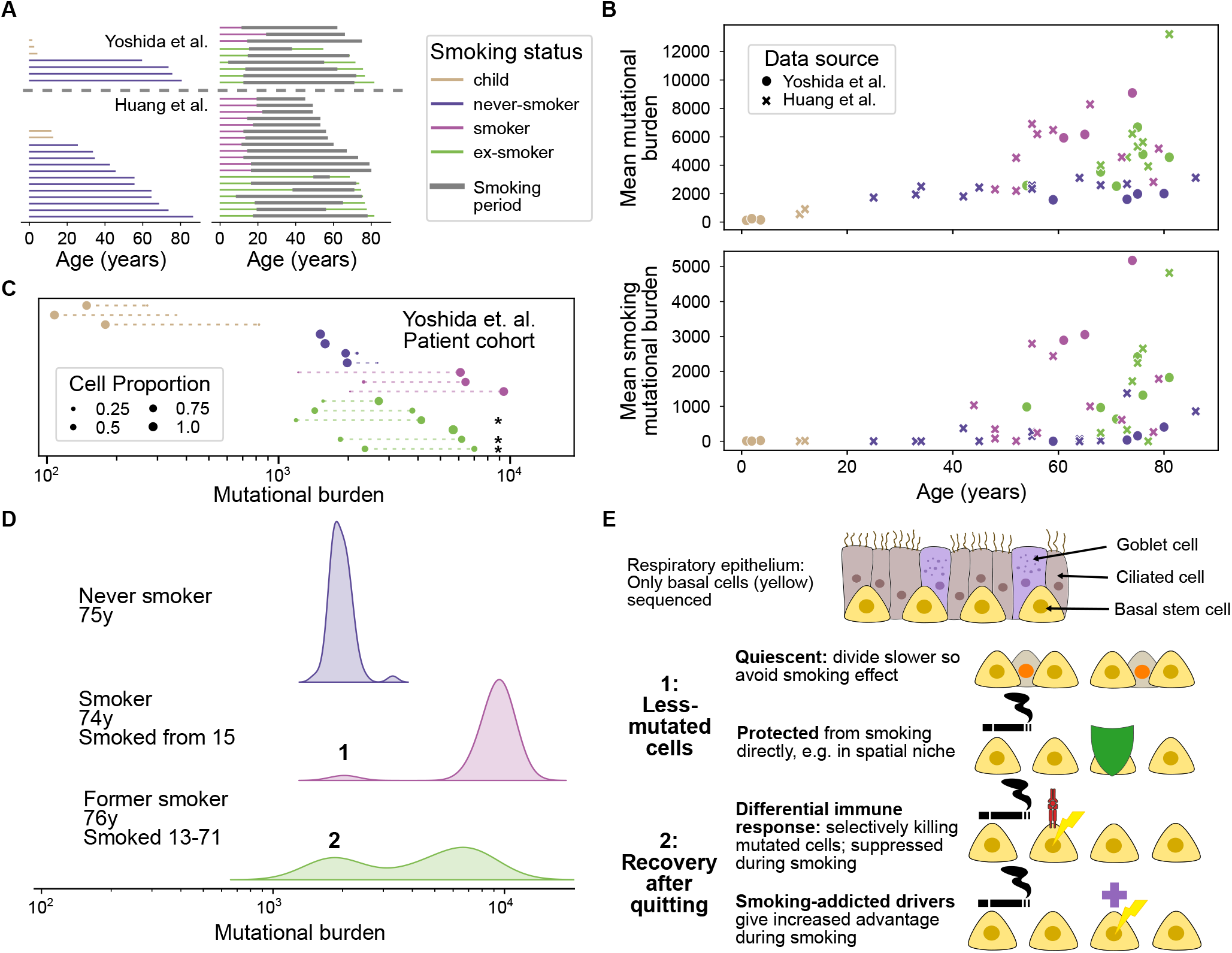
Single-cell whole-genome sequencing data from healthy stem cells shows a subpopulation less mutated by lifelong smoking and recovery of less-mutated cells after quitting smoking. (A) Smoking history distribution within the combined Yoshida et. al. (above dashed line) and Huang et. al. (below dashed line) cohorts. (B) Mean mutational burden (above) and smoking-signature mutational burden (below, corresponding to signatures SBS4, SBS92, DBS2 and ID3) across all cells sequenced from each patient, by age, smoking status and study. (C) Gaussian mixture model representations fit by expectation maximisation (see Methods), with one or two components selected by Bayesian Information Criterion (BIC), showing bimodality among ever-smokers and a recovery of the less-mutated population in ex-smokers. (*) marks significantly bimodal distributions (by each of a selection of bimodality tests with Benjamini-Hochberg correction across the cohort, see Methods). (D) Mutational burden distributions within the lungs of three patients: a 75-year-old never smoker (data from 55 cells), a 74-year-old smoker who has smoked since the age of 15 (data from 49 cells), and a 76-year-old former smoker who smoked from 13 to 71 (data from 54 cells). Kernel density estimate representations generated by the *seaborn* package^16^ using Scott’s method for bandwidth calculation^17^. (E) Resulting hypotheses for mechanisms that may bring about these unexpected dynamics.

Hypothetical explanations for these dynamics can be divided into broad classes based on the manner in which they explain each dynamic (shown in Figure 1E). To explain the bimodality, some differential reaction to smoking is needed within the stem cell population. The cells that are near-normally mutated could be daughter cells of a quiescent subpopulation of stem cells which divide much slower and hence mutate less. Such a population may exist within submucosal gland-resident myoepithelial cells, which have been shown to repopulate damaged epithelium in the event of severe injury^18^. Alternatively, the less-mutated basal cells may occupy a protected niche in airways where tobacco carcinogens cannot penetrate. These may be protected by epithelial topography or by the presence of “hillock” cells^19,20^.

To explain recovery after smoking cessation, a differential selection of a high mutational burden may be required depending on the presence or absence of tobacco smoke. This selection can either be positive or negative. Positive selection could be the result of driver mutations that confer an augmented fitness effect in the presence of smoke: a substantially altered lung environment^21^. Negative selection could come from immune surveillance that preferentially targets neoantigen-expressing highly mutated cells, but is suppressed during smoking^22^.

Limited data exists concerning the evolutionary mechanisms at work within the stem cell population in the human upper airway epithelium. Teixeira et al.^23^ used lineage tracing of naturally occurring mitochondrial DNA mutations to demonstrate a good fit to a simple theoretical model of stem cell homeostasis^24^. Lineage tracing experiments in the mouse trachea have shown an excellent fit to a simple model with a single progenitor population^25^ in the absence of severe injury^18^; similar experiments in other mouse epithelia have also inferred a single-progenitor model^26,27^. To our knowledge, there are no experimental models charting mutation dynamics under variable smoke exposure over lifetimes.

Here, we investigated the degree to which each combination of the mechanistic evolutionary hypotheses described in Figure 1E is able to explain the human sequencing data observed by Yoshida et al. and Huang et al., applying an established theoretical framework for somatic evolution in the upper airway epithelium^23,24^ to model entire human lifetimes of exposure, an analysis that would be impossible to undertake with current experimental approaches. We applied a mechanistic learning framework^28,29^, using diverse machine learning techniques to infer the modelling paradigm most closely approximating the observed data, with the distinct methodologies converging on the same resulting combination of hypotheses.

## RESULTS

### A modular evolutionary simulation model incorporates all hypotheses into an existing understanding of basal stem cell dynamics, and reproduces the broad observations of the scWGS dataset

To investigate the complex evolutionary interplay between tobacco and smoking within the combined single-cell derived WGS (scWGS) dataset^3,4^, we implemented a computational agent-based simulation framework of an established model of basal cell homeostasis^23,24^ (Figure 2A), over the lifetime of each patient in the cohort. Smoking histories were incorporated through changes to mutation and division rates. Somatic Darwinian evolution was modelled via cellular fitness, acting to bias cell fate towards or away from the basal compartment (Figure 2A), affected by mutations occurring stochastically on cell division (see Methods). Initial simulations using the established model under a well-mixed assumption displayed underdispersion of mutational burden distributions relative to observation (see Supplementary section “Non-spatial underdispersion”). Incorporation of a spatial lattice, upon which all hypotheses listed above were simulated using the Gillespie model with symmetric cell divisions tied to neighbour cell losses to maintain homeostasis (Figure 2B), increased the between-cell variation.

**Figure 2.**
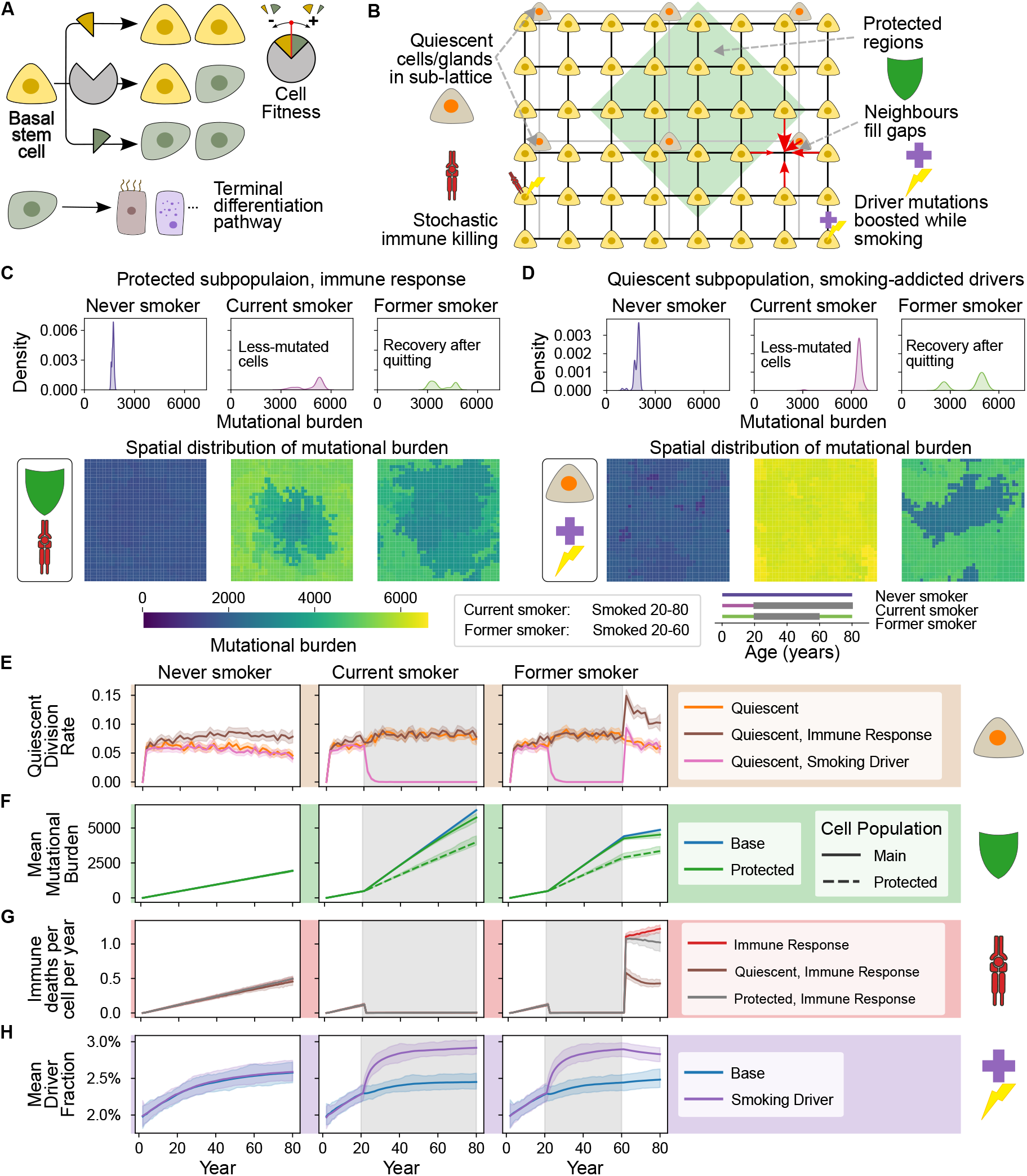
A modular evolutionary simulation model reproduces the features of observed scWGS datasets. (A) The model of lung homeostasis of Teixeira et al., adapted to include cellular fitness via fate bias. (B) Hypotheses are incorporated into a spatial lattice model of the basal membrane, using a Gillespie simulation with constant population maintained by tying together differentiations with neighbour divisions. (C) Outputs of simulations with prior mean parameter values under the Protected and Immune Response hypotheses. Distributions of mutational burden (above, kernel density estimates using Scott’s method for bandwidth calculation^17^) show similar dynamics to the observed data (Figures 1C, S1C). Spatial distribution of mutational burden on lattice (below) shows protected region less affected by smoke and clonally expanding after cessation. (D) As in Panel C, under the Quiescent and Smoking Driver hypotheses. (E-H) Demonstrations of the effect of each hypothesis on simulations with prior mean parameter values (see Supplementary section “Simulation parameters and prior distributions”), showing relevant statistics over the course of simulations. Lines show mean values in each simulated year with linear interpolation, and shaded regions show the 2.5^th^ and 97.5^th^ percentiles over 100 replicate simulations, coloured by active hypotheses. (E) Mean division rate of quiescent glands, per gland per year, binned over 2-year intervals. (F) Mean cell-location-specific mutational burden (G) Mean death rate of cells from immune killing (H) Mean fraction of mutations with positive fitness effect

The mechanistic hypotheses presented in the previous section represent four broad classes of models which could in theory explain the observations. To leverage this model for assessing the ability of each hypothesis (or combination thereof) to fit the data, minimal implementations of each were incorporated modularly such that any combination could be simulated (Figure 2B, Methods). A quiescent subpopulation is modelled as residents of glands forming a sub-lattice below the lattice of basal cells, and a protected subpopulation by regions of lattice in which the effects of smoke on mutation and division rates are reduced. Smoking-suppressed immune predation is modelled as a constant death rate of cells, proportional to the mutational burden and reduced during smoking, while smoking-addicted drivers are modelled as a linear increase in the fitness effect of driver mutations during smoking.

We first ran simulations using default parameter values (means of priors distributions derived from literature; see Methods, Table S3), to assess their general behaviour. Simulations recapitulated the broad dynamics observed by Yoshida et al. with different combinations of hypotheses (Figures 2C–2D), with bimodality in the mutational burden distribution of those with a history of smoking and a rebalancing towards less-mutated cells in a simulated ex-smoker after cessation. Examining the incorporated spatial dimension revealed localised structures emerging: cells in protected regions accumulated fewer mutations and clonally expanded after cessation (Figure 2C), whereas quiescent subpopulations led to a more evenly spatially distributed near-normally mutated population (Figure 2D). Phenomenologically, bimodality appears to arise within simulations due to a confluence of factors: negative selection of mutations gives way due to the smoking-induced selective shifts to a favouring of the more-mutated population over the less-mutated (quiescent or protected) population of cells. This allows the two populations (less fit, less-mutated and more fit, more-mutated) to co-exist until cessation of smoking, at which point the selective shift is reversed.

To confirm that simulations were modelling the evolutionary system described, we tested that each hypothesis as incorporated into the model also displayed expected interactions and effects. Each quiescent cell^1^ divides to replace basal cells at a low rate, averaging 0.53 replacements per cell/gland per year in the never-smoker. This is suppressed during smoking in the presence of the smoking-addicted driver hypothesis, as highly mutated cells have more driver mutations, and spikes after cessation of smoking if smoking-suppressed immune predation is also modelled (Figure 2E). The protected subpopulation of cells accumulates 36.5% fewer mutations during smoking, and leads to a smaller reduction of 7.9% in the mean mutational burden of the un-protected subpopulation due to clonal expansions (Figure 2F). Immune killing of healthy cells increases over the lifespan as mutations accumulate. This reaches a maximum rate of 0.48 deaths per basal cell per year in never-smokers and current smokers, significantly below the known turnover of cells in these groups^23^. This dynamic is suppressed by smoking (Figure 2G). Driver mutations are selected for over simulated decades, with an increased selection when their fitness is augmented by smoking (Figure 2H).

Combinations of hypotheses demonstrate synergistic effects, such as quiescent populations and differential immune response together reducing post-smoking mutational burden accumulation significantly more (reduced by 49.0 mutations per year) than either individually (19.0 and 0.22 mutations per year respectively; see Figure S4A).

Together, these results establish that the proposed hypotheses can qualitatively explain the dynamics observed in the combined cohort of scWGS data and confirm that the simulation framework created here produced realistic results.

### Combinations of hypotheses can be accurately inferred from unseen simulated datasets

Previous studies of mutant cell dynamics have placed emphasis on the statistical modelling of clone fate mapping data^23^. However, here, the connection between mutational burden and the statistical cellular framework is remote, calling for an alternative strategy. We generated a dataset of simulated cohorts with distinct combinations of hypotheses activated, and parameter values drawn from literature-derived prior distributions (see Supplementary section “Simulation parameters and prior distributions” and Table S3), and applied machine learning classification approaches to the inverse problem mapping a simulated dataset to the combination of hypotheses that created it. These classifiers fill the dual role of assessing identifiability and drawing inference from the observed scWGS data (Figure 3A). In order to use only relevant information, classifiers were trained on multidimensional scaling embeddings (see Methods) of simulated cohorts with respect to biologically motivated distance functions using modalities that can be extracted from scWGS data (Figure 3C).

**Figure 3.**
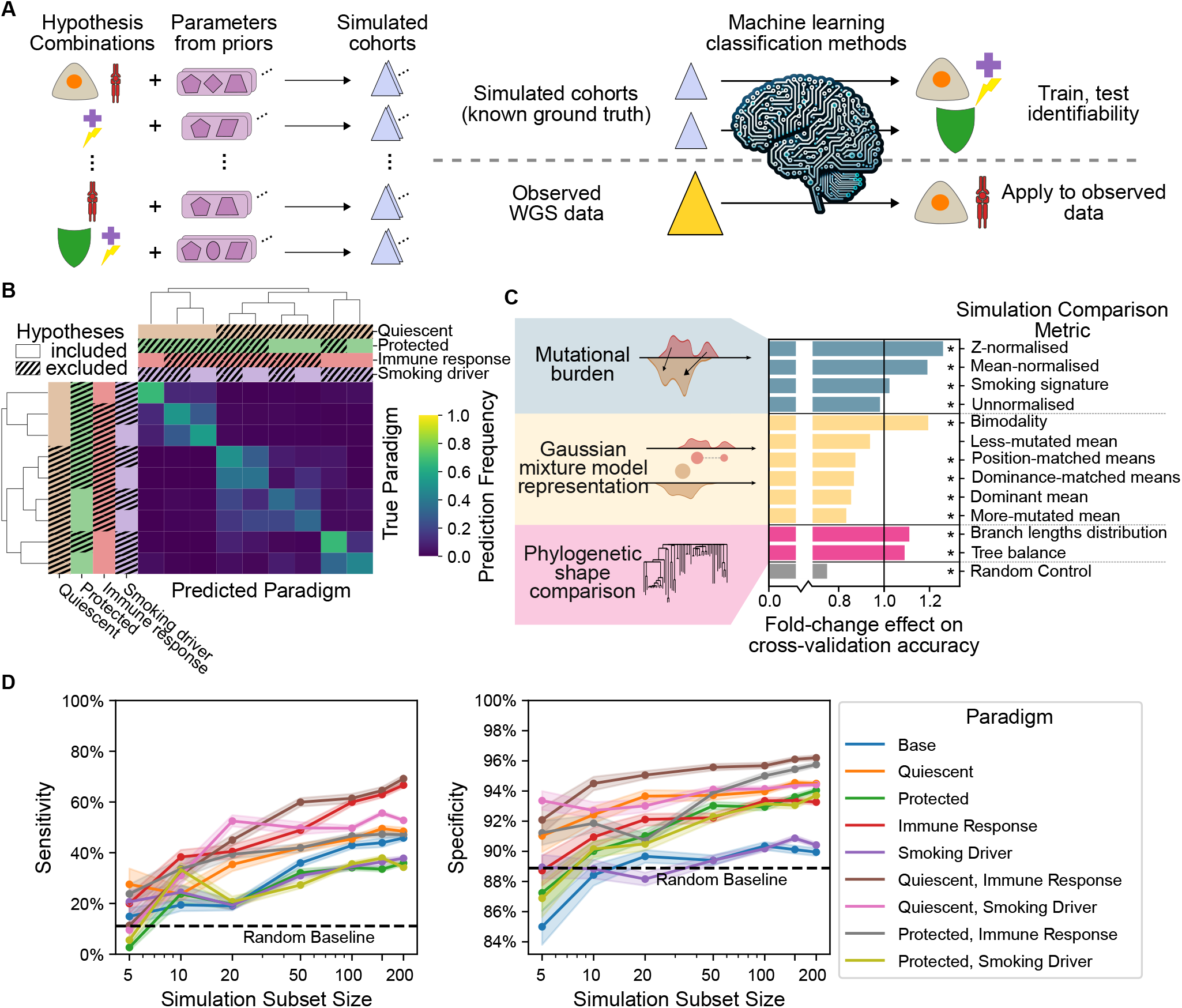
Machine learning classifiers are able to discern driving hypotheses in unseen simulated datasets. (A) Simulations are run with each combination of hypotheses, with parameter values randomly drawn from prior distributions. Classifiers are trained to find the combination of hypotheses from outputs, in order to assess the degree of identifiability and gain mechanistic inference from the observed dataset. (B) Mean classification frequency across all classifiers of each paradigm, in assessing outputs from unseen simulations with each other ‘True’ paradigm. All folds, embedding space dimensionalities, classification methodologies and technical replicates are included, for a total of 150 classifications of each simulation. (C) Impact on accuracy of each biologically-motivated metric used to train models on simulation outputs. (*) marks statistically significant effect on accuracy, by Holm-corrected t-test adjusting for feature dimensionality and classification method. Subsets of at most 3 distance functions considered (see Methods, Tables S4–S6). (D) Sensitivity (left, probability of classifying the hypothesis as present if true) and specificity (right, probability of classifying the hypothesis as absent if true) of each hypothesis, for classifiers trained on different subsamples of simulations. Points represent means, lines show interpolation, and shading shows a bootstrapped 95% CI over 3 random subsamples of training simulations at each size.

While classifiers showed limited raw accuracy (52.9% for random forest and 53.6% for logistic regression, Figure S9A), the misclassifications were not random: confusion matrices (Figures 3B, S9B) reveal a structure of subgroups of paradigms whose simulation outputs were highly distinguishable. Hierarchical clustering of these same matrices reveals which hypotheses left a detectable mark in scWGS data: the effect of the smoking-addicted drivers hypothesis was less detectable in all combinations, while the quiescent hypothesis left a clear signature.

Next, we assessed which metrics were used by the classifiers in making inferences on the mechanistic hypotheses active in generating a particular simulated cohort. For the classification methodologies used here, there are internal importance scores that can be extracted for each distance function (Figures S10A–S10B), but these have limited accuracy. For a reliable indication we instead compared the accuracy of classifiers trained with and without each metric (see Methods). By this measure, classifiers principally used normalised comparisons of mutational burden distributions (providing a 21.6% increase in accuracy for Z-normalised and a 15.9% increase for mean-normalised), the degree of bimodality of the same distribution (16.2% increase), and phylogenetic shape metrics (9.26% increase for branch lengths distribution, 7.52% for tree balance) in their inference (Figure 3C).

A common problem with using machine learning to infer the invertibility of a mapping is accidental labelling^30^. In order to assess whether the degree of accuracy reached here is due to this issue, we retrained classifiers on randomly subsampled simulation sets. These subsampled simulations showed reduced sensitivity and specificity, reaching near-random levels at 5 training simulations per hypothesis combination. This would be unlikely if previous results were results of a strong spurious labelling effect.

Overall, this mechanistic analysis demonstrates that the scWGS data present in the cohort is sufficient to distinguish which combination of hypotheses is most likely to occur in the human lung.

### Classifiers predict a quiescent subpopulation and differential immune response as key mechanisms explaining observed data

The classifiers trained above map a simulated dataset to the known combination of hypotheses active in its creation. We next applied these classifiers to ascertain which mechanistic hypotheses were at work in the human lungs of the cohorts of Yoshida et al and Huang et al^3,4^. We embedded these observed scWGS datasets via the same procedure and distance metrics (Figure 3A).

To ensure the accuracy shown in Figure 3B was maintained on a new unseen datapoint, distance functions which placed the observed dataset outside the distribution of the simulations were excluded, at multiple thresholds for extremity (Figures 4A–4B, Methods).

**Figure 4.**
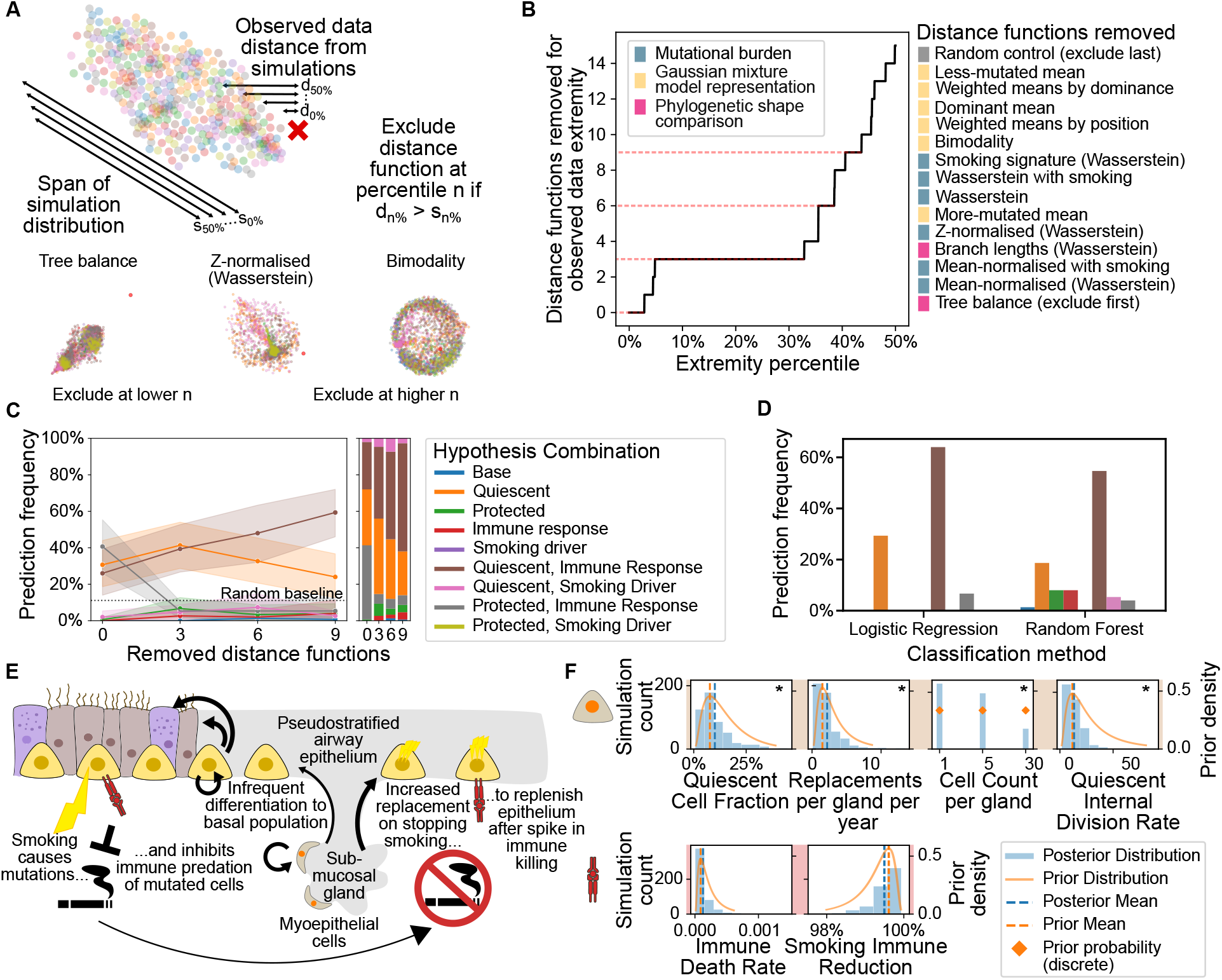
Applying classifiers to observed scWGS datasets reveals a quiescent subpopulation of cells and smoking-induced suppression of immune predation of healthy lung basal stem cells as key drivers of the observed dynamics. (A) Distance functions are filtered by the extremity of the true data within the distribution of simulations. Examples of MDS projections (see Methods, Figure S8) of three distance functions (Tables S4–S6) are shown for illustration, removed at thresholds 2.8%, 35.5%, and 43.6% respectively. (B) Number of distance functions excluded after filtration with different thresholds of observed data extremity. Thresholds chosen for the remainder of the work are shown in red; 2-dimensional MDS projections are shown in order of elimination in Figure S8; metrics are described in Tables S4–S6. (C) Frequency of classifying the observed dataset within each paradigm, by true data extremity threshold, showing selection of the quiescent and immune response hypotheses at the most stringent observed data extremity threshold. Points show mean frequency over all predictions (over 5 folds, 2 classification methods, 3 embedding replicates, 5 feature space dimensionalities for a total of 150 predictions at each threshold). Lines show interpolation, and shaded area represents a bootstrap 95% CI over technical replicate classifiers. A baseline of random selection of one out of 9 paradigms is shown for comparison. (D) At the most stringent threshold for observed data extremity, the frequency of classification as each paradigm among all replicate classifiers of the true data, split by classification methodology. (E) Proposed resulting paradigm of lung evolution in response to smoking and smoking cessation: sub-mucosal gland-resident myoepithelial cells replenish the airway after a spike in immune killing on stopping smoking. Further validation is required to discern details with certainty. (F) Distribution of parameters specifying the quiescent subpopulation and immune predation in 5% of simulations most closely approximating (by summed normalised distances) the observed dataset, within the quiescent and immune response paradigm, at the most stringent observed data extremity threshold. 500 simulations shown out of 10,000 run with parameters drawn from prior distributions (shown in yellow, as probability density or mass functions). Prior and posterior means are shown. Asterisks denote parameters for which the posterior distribution is significantly different to the prior distribution (see Table S8 for p-values and a full description of the tests).

Technical replicates of the embedding procedure and folds in 5-fold cross-validation provided a distribution of classification predictions for each level of observed data extremity filtering. Without filtering, this distribution was split over multiple possible paradigms (likely reflecting quasi-random assignment to an out-of-training-distribution data-point), but with filtering classifiers indicated a quiescent subpopulation of cells combined with differential immune predation as the likeliest paradigm explaining the observed scWGS dataset in 59.3% of predictions (Figure 4C).

This result was consistent across distinct classification methodologies (Figure 4D). Furthermore, an alternative route of inference via aggregating distance functions concurred (Figures S12B–S12C), indicating that this result was unlikely to be an artefact of the embedding procedure. The resulting proposed paradigm is illustrated in Figure 4E. Interrogating model parameters enables us to provide a quantitative understanding of these dynamics (Figure 4F), and infers a small number of quiescent cells in each gland, with quiescent cells comprising 10.1% of the stem cell pool, as well as a mutation rate of 46.1 mutations per cell per year (Figure S11D), which is above the prior estimate (see Supplementary section “Simulation parameters and prior distributions” and Table S3).

## DISCUSSION

Despite decades of research establishing a clear link between tobacco smoking and lung cancer risk, mechanistic understanding of the risk dynamics associated with quitting smoking has remained stubbornly limited. Here, we used patterns of somatic genetic damage to establish a paradigm for explaining the remarkable recovery in genomic damage and cancer risk, with smoking cessation re-establishing immune predation of highly mutated stem cells and thus allowing slow-cycling stem cells to repopulate the airway.

First, we created an *in silico* model for the somatic evolution that occurs in healthy upper airway epithelial tissue over an entire lifetime, using smoking histories to individualise the model to each patient’s life course. The framework incorporates mechanistic hypotheses modularly, and demonstrates a qualitative fit to the observed recovery in basal cell mutational burden after smoking cessation. This validates previous speculations^3^ as to the mechanisms behind the puzzling data.

We then created a dataset of simulated cohorts to assess the distribution of simulation outputs. Using machine learning classification methods, we demonstrated that despite all fitting the observed dynamics qualitatively, the differing simulations were distinct quantitatively, distinguishable from outputs alone.

Finally, we applied classifiers trained on simulations to the observed scWGS dataset, and ascertained using orthogonal approaches that the most likely paradigm at work in the human lung is a slow-cycling population of stem cells replenishing the lung after tobacco smoke’s suppression of immune predation is removed. This was supported by quantitative analysis despite other combinations showing a qualitative fit.

This work could not have been carried out using current experimental protocols *in vivo*: in the inference procedures used, almost 70 million patient-years were simulated (see Supplementary section “Total patient-years simulated”). This is inherent to the problem of studying lifelong somatic evolutionary dynamics, which occur over long timescales and with a high level of stochasticity.

The specific distance metrics used for this inference, those not removed for placing the observed data outside the distribution of simulations, were principally concerned with Gaussian mixture models fit to the data, suggesting a coarse-grained view on the data as the correct level of resolution, with good fit to the data and sufficient resolution to distinguish between hypothetical models.

While this study provides a first attempt to mechanistically model somatic evolution in response to smoking at the scale of a lifetime, there remain limitations to our work: small patient cohorts and consequent limited personalisation of simulations mean that interpersonal heterogeneity is not considered in full, with focus rather on general evolutionary dynamics. Opting for an agent-based modelling system permits more nuanced interrogation and understanding of the underlying complexity than commonly used alternatives such as systems of differential equations, but at the cost of increased reliance on validity of prior parameter distributions and increased computational complexity (limiting inference methodologies). Nonetheless, as with much work in mathematical modelling of biological systems, model parsimony has been favoured over complexity in order to compare only the minimal examples of each of the four classes of model. This may limit the precision of the resulting paradigm: for instance, a more involved model of immunity could require a larger mutated clone to trigger a reaction, or model exhaustion with sustained large mutational load. Finally, although we inferred our *in silico* results in two independent cohorts, interventional studies *in vitro* or *in vivo* would provide additional confidence in our proposed paradigm (Figure 4E) and guide the next experimental steps.

Indeed, this work provides a strong direction for such future research: the signals found here suggest a paradigm of evolution which would equally leave its mark in other modalities, such as co-evolution of lung and immune cells or expression of transcriptomic signatures associated with submucosal gland-resident cells in the local epithelium after smoking cessation. Linking this work to established epidemiological trends (via simple assumptions around mutational burden and risk of cancer initiation) could forge a direct link between a previously undetected tissue-level phenomenon and its signature at a population scale. This model may further be re-used for *in silico* trials of future preventative lung cancer therapeutics, providing a rapid, perturbable system modelling accumulation of genomic damage in the presence of carcinogens and immune modulation, an invaluable resource for researchers aiming to reduce the burden of disease on society.

## METHODS

### Data re-analysis

While the majority of data processing was completed as part of the two papers from which this work draws data^3,4^, some additional analysis was required. Children were defined as those under the age of 5 years.

#### Mutational signature decomposition

In order to incorporate newly discovered smoking-associated mutational signature SBS92^7^, as well as to ensure consistency of analysis methods, signature alignment performed in the source publications were re-calculated. All mutations were re-assigned to mutational signatures using the SigProfilerAssignment tool (Version 0.1.0^31^). Single base substitutions (SBS), double base substitutions (DBS) and insertion-deletion (ID) signatures were assigned separately to COSMIC signatures^32^ occurring in LUSC or LUAD samples in the Pan-Cancer Analysis of Whole Genomes (PCAWG) cohort^33^, as well as newly discovered signature SBS92 which was shown to occur in LUSC in the Tracking Cancer Evolution through Therapy (TRACERx) cohort^34^. SBS8 was excluded as occurring principally after tumourigenesis^35^. As in the analysis by Yoshida et al.^3^, the majority of mutations were assigned to clock-like signature SBS5 (Figure S2A), while smokers and ex-smokers exhibited significantly higher proportions of smoking signatures SBS4, SBS92, DBS2 and ID3 (Figures S2B–S2C).

This re-analysis provided a separation of mutational burden into smoking-attributable mutations (those assigned to SBS4^36^, SBS92^7^, DBS2^37^ or ID3^33^) and other mutations (Figure S1B).

#### Mutational burden bimodality testing

Distributions of mutational burden for each patient were tested for bimodality against 6 different bimodality tests^38–43^, as implemented in the *multimode* R package^43^ using a bootstrap replicate value of 10^5^. All tests concurred in assigning significance to those patients marked with (*) in Figure 1C (p-values given in Table S1).

#### Gaussian mixture model fitting

To fit 1- or 2-component Gaussian mixture models, an expectation maximisation algorithm was used as implemented by the *scikit-learn* Python package^44^ for each component count. These two models were then compared for BIC, and the model with a lower value was chosen.

#### Phylogenetic tree summary

Phylogenies were generated by Yoshida et al., but not by Huang et al. due to reduced cell counts per patient (Figure S1A). In order to compare these phylogenies with simulated phylogenies, summaries were required that capture the evolutionary dynamics. Two summaries were chosen: the distributions of branch lengths, which showed visible variation by smoking status (Figure S2D), and the tree balance metric J_1_^45^ Figures S2E–S2F).

### Simulation model high-level summary

The model used to run simulations, described in detail in the Supplementary Text (section “Modelling framework”), is an agent-based model in which each cell follows the division schema shown in Figure 2A^23,24^. Cells divide stochastically at a constant rate via a Gillespie simulation^46^, and accumulate a stochastic number of mutations with each division. Some fraction of mutations bring a fitness change, which subsequently changes the cell’s division type distribution (Figure 2A).

This model is initialised on a 40 *×* 40 spatial lattice (far larger than the average clone size after 60 years in simplified simulations; see Figures S5A–S5B) for each patient in the combined cohort, and simulated for the duration of that patient’s lifetime up to the age at which the sample was taken. If at a specific age the patient was smoking, the mutation rate and division rate are increased. The edges of the lattice are joined together to form a torus to remove edge effects. When cells are removed, either by differentiation or immune death, neighbours compete to fill the gap according to their symmetric division rate, thus maintaining constant population size by tying together removal of stem cells with production of new cells.

The hypotheses described in the introduction are included modularly, such that any combination can be activated for any given simulation of the cohort. Combinations of activated hypotheses are referred to as hypothetical paradigms. Hypotheses are incorporated as follows:

- Quiescent cells are modelled as residents of glands containing one or more stem cell, in which fixation of new mutations is instant and stochastically dependent on the fitness of the new mutation. These glands form a sub-lattice below the lattice of basal cells, and compete to fill gaps only in the cell directly above them.
- The protected subpopulation is modelled as regions of the lattice in which the effect of smoke on mutation and division rates are linearly reduced.
- Smoking-suppressed immune predation is modelled as a constant death rate of cells, proportional to the number of mutations they have accumulated.
- Smoking-addicted drivers are modelled as a multiplicative increase applied to the fitness effect of every mutation with positive fitness effect, which applies for the duration of smoking for mutations acquired before or during smoking.

To draw samples for comparison with other simulations or the observed scWGS datset, a random subsample is drawn from the population of cells at the age of sampling (mimicking the data gathering procedure; see Supplementary section “Sampling from simulations”). Phylogenies are reconstructed, ignoring all cells not in the subsample.

### Biologically motivated simulation comparisons

The outputs that may be extracted from the simulations used in this work have many dimensions and are stochastic. Because of this, the likelihood of exactly reproducing any given dataset (such as we may hope to do with the observed scWGS dataset) is negligible.

In order to make inferences using our simulation framework, metrics were required to capture relevant characteristics of simulation outputs, while only using information that could be gathered from the data in question. In this instance, the data with which simulation outputs can be compared is single-cell derived WGS data from multiple healthy lung basal cells from the same patient^3,4^. This data can yield:

- Counts of somatic mutations incurred by each cell over the course of the individual’s lifetime;
- Counts of mutations attributable to each known mutational signature (by trinucleotide context^47^), in particular those signatures whose aetiologies are known to be related to tobacco smoking (see above);
- Features of phylogenetic trees inferred from shared mutations between cells (only for those from the Yoshida et al. cohort, as the number of cells sequenced per patient in the Huang et al. cohort is too low for meaningful phylogeny comparison Figure S1A).

Many forms of information often reported from WGS data, such as specific driver mutations, were not applicable in this case: simulations track only generic, unnamed mutations in order to have reasonable likelihood of replicating the shape of the observed data. Due to the relatively low cell counts from each patient, variant allele frequencies would be low resolution and are thus omitted.

The metrics referred to here include, but are not limited to, those based on “summary statistics” of the data. These can be useful for visualisation and for clarifying what exactly is relevant about a dataset, and are commonly used in simulation studies (particularly those using Bayesian approaches^48^). However, the concept of comparing two datasets need not be restricted to those metrics that can be expressed as a composition of a projection into a Euclidean space followed by application of its canonical metric. By avoiding this restriction, it was possible to consider more varied metrics such as the 1-Wasserstein metric^49^.

#### Wasserstein distance of mutational burden distribution

The principal surprising results from the Yoshida et al. paper concern the distribution of mutational burden across the cells of the healthy human lung (namely its approximate bimodality in those with a history of smoking and the increase in the fraction of cells with near-normal mutational burdens in ex-smokers compared to current smokers; see Introduction). As such, a prime candidate for a relevant comparison between cohorts is to assess how closely aligned their distributions of mutational burden are.

Here, presented with two empirical distributions D_obs_ and D_sim_ and faced with the question “how likely is it that D_obs_ was created by sampling from D_sim_?”, a natural conclusion would be to use the Kullbach-Liebler divergence, which has the useful property of directly answering this question^50^. However, this metric is inappropriate in sparse distributions such as these single-cell empirical mutational burden distributions, due to its problems with 0 values in the source distribution (D_sim_ in this case). Instead of adding complexity to work around this issue, say using a kernel density estimate to smooth out stochastically occurring zero values in areas of non-zero support, we used the 1-Wasserstein metric^49^. This distance function measures the minimal total work required to move one distribution’s probability density to another’s, earning it the alternate name the Earth-mover distance. The mathematical definition of the Wasserstein distance is given above; the Python implementation in *scipy*.*stats*.*wasserstein_distance* is used here^51^.

Noting that the 1-Wasserstein distance between distributions of similar shapes but different mean values can be considerably higher than that if the distributions had different shapes but the same mean, another possibly useful comparison given the importance of the shapes of the distribution functions is the 1-Wasserstein distance after first normalising the data. We therefore also created mean-subtracted and z-transformed metrics given by

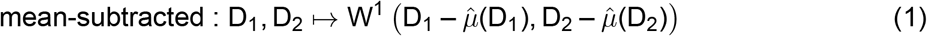

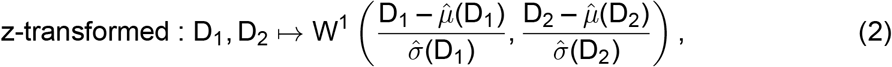

where 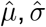 are the sample mean and standard deviation respectively, and D_1_, D_2_ are sets of non-negative integers (or equivalently the empirical distribution they generate) representing single-cell mutational burden distributions.

#### Gaussian mixture model-based distances

As described above, a natural way to summarise distributions of mutational burden in this dataset is to fit a one- or two-component Gaussian mixture model. This then provides another means of comparison, directly comparing the parameters of the fitted mixture models of the two distributions, which gives rise to multiple possible metrics, each of which can be considered in either absolute value or squared value for each patient:

- Difference between the larger weight of the two models
- Difference between the means of the models’ dominant or subdominant peaks (the component with the larger weight or smaller weight, respectively)
- Difference between the means of the models’ higher or lower peaks (the component with a larger or smaller mean, respectively)
- Weighted sums of squared comparisons of the means, weighted by the product of the weights of two compared components; components can be matched by either dominance or position.

As the spatial simulation implementation only partially reduced the issue of underdispersion of mutational burden observed in non-spatial simulations (particularly within never-smokers, see Supplementary section “Non-spatial underdispersion”) these metrics all ignore the standard deviations of the two peaks so as not to simply choose the simulation which most drastically increases its within-subpopulation variance.

#### Smoking signature mutational burden

As described above, the trinucleotide context of sequenced mutations can be used to decompose the count of mutations in each cell into smoking-attributable mutations and others. This can also be tracked within simulations (above). This new distribution of smoking-attributable mutation across cells can again be compared using the same 1-Wasserstein distance to compare two cohorts.

This decomposition also provides a natural two-dimensional representation of each cell by its number of smoking-attributable and non-smoking-attributable mutations (as in Figure 1B). A two-dimensional 1-Wasserstein distance, achieved using the *linear_sum_assignment* function from the *scipy*.*optimize* package, can be used to calculate a metric which fully considers the two distributions’ dissimilarity. The high computational cost of this calculation leads it to be infeasible in large sets of simulated cohorts; to check whether the combination was likely to be a major omission a “simplified” version of the distance was included by simply summing the two marginal 1-dimension 1-Wasserstein distances. Analyses showed this approximation did not contribute significantly therefore the true 2-dimensional distance was not calculated at scale.

#### Phylogenetic tree comparison

Phylogenies generated from simulations can be compared with the observed phylogenies via summaries extracted from each tree, as described above: the J^1^ tree balance index, and the distribution of branch lengths. As these values may be sensitive to subsampling of the phylogenetic tree, this is run on a randomly selected subsample of the final population of cells (see above). For the tree balance index, either the absolute or squared difference can be taken for each patient, and the branch length distribution can again be compared using the 1-Wasserstein distance. As an additional possible source of information, the branch lengths can be summed and compared by absolute value; this sum will have higher values if cells are less related.

#### Negative control distances

To bolster against spurious results from any inference procedures based on these distance metrics, we included two distances which include no information from the two cohorts being compared: one which provides a random number between 0 and 1 for each patient, and one providing the number 0 for every comparison. These additionally provide a baseline for any significance measures: any level of significance assigned to these is from random noise and can be discounted.

#### Aggregation over patient cohort

Each simulation is run for each patient separately, taking into account their smoking history. Each comparison metric compares two simulated cohorts patient-wise: the overall value for the distances between the two cohorts is taken to be the weighted sum of the comparison for each patient, weighted by the log-scaled number of cells sequenced from that patient. The rationale for this weighting is that more information should be drawn from patients for whom there is more sequenced data, and the effective sample size shouldn’t be inflated when comparing simulated cohorts with each other even if there are a larger number of cells to theoretically subsample from. The log-scale is used due to heuristics from information theory^52^: an additional unit of information is gained by doubling the number of samples, rather than by adding a fixed number of samples.

### Identifiability analysis

To determine identifiability of simulation outputs, we generated an equal number of simulations in each of 24 paradigms. We calculated all pairwise comparisons for all pairs of simulations, and used these distances to embed each simulation into a separate n-dimensional space for each distance function using multi-dimensional scaling, with n taking values 2, 3, 5, 10 and 20 We then concatenated the MDS embeddings of each distance function to form a single vector for each simulation, and trained machine learning classifiers (logistic regression, support vector machine and random forest methodologies) to predict the combination of hypotheses generating a simulation from its concatenated MDS embedding. These classifiers were tested for accuracy via 5-fold cross-validation.

We initially created a dataset of 100 simulations per paradigm, with two replicates of each simulation. Due to the low between-replicate distances observed, we created and trained classifiers on a larger dataset of 300 simulations per paradigm with only one replicate. Based on lower accuracy when trained on this dataset (Figure S9A), the support vector machine methodology was excluded for downstream analysis.

An internal importance score of each feature in training classifiers can be calculated:

- In a logistic regression classifier, importance of each feature can be inferred by normalising features before training the classifier and taking the absolute value of the coefficient assigned to each feature.
- In a random forest classifier, the Gini importance value is used^53^.

The mean value of the importance scores across the features contributed by a particular distance function are taken as the raw importance score for that metric (Figure S10A). As these scores are not always viewed as accurate^53^, a more reliable measure was created by re-running the training using MDS embeddings of only subsets of distance functions. This was run for all subsets of at most three distance functions, or at least 14 out of 17 total. The fold change in accuracy between simulations with and without each distance function were used to assess their importance in training, separately in the larger and smaller subsets of distance functions (Figure S10B).

In addition to restricting the distance functions on which classifiers were trained, we also restricted the number of simulations’ embeddings used to train classifiers (Figures S10C–S10D) and the set of patients over whom the distance function was calculated (Figure S10E).

To apply to the observed scWGS dataset, distances from this data to each simulation was calculated, and included in the MDS embedding. Distance functions with respect to which the true data was outside the distribution of simulations were excluded as described in the main text and Figure 4A. Classifiers trained on the remaining subset of distance functions at each threshold were applied to predict the paradigm of the observed dataset (Figures 4C– 4D). Additional classifiers were trained on subsets of simulations, as well as these subsets of distance functions (Figure S12A).

We subsequently created a dataset of 10,000 simulations in the quiescent and immune response paradigm, and calculated only the distances to the observed dataset. We then ranked these simulations by the aggregated distance (see Supplementary section “Distance function aggregation”) to the observed dataset; the parameter values of the 5% closest simulations are shown in Figure 4F and Figure S11D. Statistical testing showed significant differences between prior and posterior distributions in nearly all parameters (Table S8).

## Data and Code Availability

The scWGS datasets are available in processed form via the original publications^3,4^.

All code used to run simulations, inference pipelines and visualise data are available at https://github.com/hughselway/SmokingSomaticEvolutionModel.

## Acknowledgements

This work was supported by MRC Programme grant MR/W025051/1 (S.M.J.); CRUK Programme award EDDCPGM\100002 (S.M.J.); UK Engineering and Physical Sciences Research Council EP/P034616/1 (B.D.S.); Wellcome Trust 219478/Z/19/Z (B.D.S.); EP Abraham Research Professorship RP/R1/180165 (B.D.S.); CRUK grant CDEPIL-Jan24/100032 (B.A.H.); CRUK Lung Cancer Centre of Excellence C11496/A30025 (S.M.J.); CRUK City of London Centre (S.M.J.); University College London Hospitals Charitable Foundation, National Institute for Health Research (NIHR) University College London Hospitals Biomedical Research Centre (S.M.J. and A.P.). N.M. receives funding from Cancer Research UK (CRUK) (DRCPFA-Nov23/100003) and has received funding from the Wellcome Trust and the Royal Society (211179/Z/18/Z) relevant to this work. N.M. also receives funding from Cancer Research UK Lung Cancer Centre of Excellence, Rosetrees, and the NIHR BRC at University College London Hospitals. C.M.-R. is supported by CRUK. B.A.H. acknowledges support from the Royal Society (Grant No. UF130039).

The authors are grateful to past and present members of the Janes lab for ongoing feedback and support, and to Helena Coggan and James Bayliss for valuable discussions during the conceptualisation and delivery of this project.

For the purpose of Open Access, the authors have applied a CC BY public copyright licence to any Author Accepted Manuscript version arising from this submission.

## Author Contributions

Hugh Selway-Clarke: Conceptualisation, Formal analysis, Investigation, Methodology, Software, Visualisation, Writing - original draft. Adam Pennycuick: Conceptualisation, Methodology, Supervision, Writing - review and editing. Sam M. Janes: Conceptualisation, Funding acquisition, Resources. Anob Chakrabarti: Conceptualisation, Methodology, Supervision, Writing - review and editing. Kate Gowers, Benjamin D. Simons, Benjamin A. Hall, Calum Gabbutt: Conceptualisation, Supervision. Nicholas McGranahan: Conceptualisation, Resources, Supervision. Ahmed Alhendi, Carlos Martínez-Ruiz, Vitor H. Teixeira: Supervision.

## Footnotes

1 A “quiescent cell” represents either a single cell or a gland containing multiple cells, depending on a parameter of the model.

## References

1. Kenfield, S. A., Wei, E. K., Stampfer, M. J., Rosner, B. A., and Colditz, G. A. (2008). Comparison of Aspects of Smoking Among Four Histologic Types of Lung Cancer. Tobacco control 17, 198–204.

2. Hecht, S. S. (2008). Progress and Challenges in Selected Areas of Tobacco Carcinogenesis. Chemical Research in Toxicology 21, 160–171.

3. Yoshida, K., Gowers, K. H. C., Lee-Six, H., Chandrasekharan, D. P., Coorens, T., Maughan, E. F., Beal, K., Menzies, A., Millar, F. R., Anderson, E., et al. (2020). Tobacco smoking and somatic mutations in human bronchial epithelium. Nature 578, 266–272.

4. Huang, Z., Sun, S., Lee, M., Maslov, A. Y., Shi, M., Waldman, S., Marsh, A., Siddiqui, T., Dong, X., Peter, Y., et al. (2022). Single-cell analysis of somatic mutations in human bronchial epithelial cells in relation to aging and smoking. Nature Genetics 54, 492–498.

5. Martincorena, I. (2019). Somatic mutation and clonal expansions in human tissues. Genome Medicine 11, 35.

6. Brunner, S. F., Roberts, N. D., Wylie, L. A., Moore, L., Aitken, S. J., Davies, S. E., Sanders, M. A., Ellis, P., Alder, C., Hooks, Y., et al. (2019). Somatic mutations and clonal dynamics in healthy and cirrhotic human liver. Nature 574, 538–542.

7. Lawson, A. R. J., Abascal, F., Coorens, T. H. H., Hooks, Y., O’Neill, L., Latimer, C., Raine, K., Sanders, M. A., Warren, A. Y., Mahbubani, K. T. A., et al. (2020). Extensive heterogeneity in somatic mutation and selection in the human bladder. Science 370, 75–82.

8. Martincorena, I., Fowler, J. C., Wabik, A., Lawson, A. R. J., Abascal, F., Hall, M. W. J., Cagan, A., Murai, K., Mahbubani, K., Stratton, M. R., et al. (2018). Somatic mutant clones colonize the human esophagus with age. Science 362, 911–917.

9. Martincorena, I., Roshan, A., Gerstung, M., Ellis, P., Van Loo, P., McLaren, S., Wedge, D. C., Fullam, A., Alexandrov, L. B., Tubio, J. M., et al. (2015). Tumor evolution. High burden and pervasive positive selection of somatic mutations in normal human skin. Science (New York, N.Y.) 348, 880–886.

10. Moore, L., Cagan, A., Coorens, T. H. H., Neville, M. D. C., Sanghvi, R., Sanders, M. A., Oliver, T. R. W., Leongamornlert, D., Ellis, P., Noorani, A., et al. (2021). The mutational landscape of human somatic and germline cells. Nature 597, 381–386.

11. Mitchell, E., Spencer Chapman, M., Williams, N., Dawson, K. J., Mende, N., Calderbank, E. F., Jung, H., Mitchell, T., Coorens, T. H. H., Spencer, D. H., et al. (2022). Clonal dynamics of haematopoiesis across the human lifespan. Nature 606, 343–350.

12. Lee-Six, H., Olafsson, S., Ellis, P., Osborne, R. J., Sanders, M. A., Moore, L., Georgakopoulos, N., Torrente, F., Noorani, A., Goddard, M., et al. (2019). The landscape of somatic mutation in normal colorectal epithelial cells. Nature 574, 532–537.

13. Hill, W., Lim, E. L., Weeden, C. E., Lee, C., Augustine, M., Chen, K., Kuan, F.-C., Marongiu, F., Evans, E. J., Moore, D. A., et al. (2023). Lung adenocarcinoma promotion by air pollutants. Nature 616, 159–167.

14. Barrett, J. C. (1993). Mechanisms of multistep carcinogenesis and carcinogen risk assessment. Environmental Health Perspectives 100, 9.

15. Lawson, A. R. J., Abascal, F., Nicola, P. A., Lensing, S. V., Roberts, A. L., Kalantzis, G., Baez-Ortega, A., Brzozowska, N., Moustafa, J. S. E.-S., Vaitkute, D., et al. (2024). Somatic mutation and selection at epidemiological scale.

16. Waskom, M. L. (2021). seaborn: statistical data visualization. Journal of Open Source Software 6, 3021.

17. Scott, D. W. (1979). On optimal and data-based histograms. Biometrika 66, 605–610.

18. Lynch, T. J., Anderson, P. J., Rotti, P. G., Tyler, S. R., Crooke, A. K., Choi, S. H., Montoro, D. T., Silverman, C. L., Shahin, W., Zhao, R., et al. (2018). Submucosal Gland Myoepithelial Cells Are Reserve Stem Cells That Can Regenerate Mouse Tracheal Epithelium. Cell Stem Cell 22, 653–667.e5.

19. Montoro, D. T., Haber, A. L., Biton, M., Vinarsky, V., Lin, B., Birket, S. E., Yuan, F., Chen, S., Leung, H. M., Villoria, J., et al. (2018). A revised airway epithelial hierarchy includes CFTR-expressing ionocytes. Nature 560, 319–324.

20. Lin, B., Shah, V. S., Chernoff, C., Sun, J., Shipkovenska, G. G., Vinarsky, V., Waghray, A., Xu, J., Leduc, A. D., Hintschich, C. A., et al. (2024). Airway hillocks are injury-resistant reservoirs of unique plastic stem cells. Nature 629, 869–877.

21. Duclos, G. E., Teixeira, V. H., Autissier, P., Gesthalter, Y. B., Reinders-Luinge, M. A., Terrano, R., Dumas, Y. M., Liu, G., Mazzilli, S. A., Brandsma, C.-A., et al. (2019). Characterizing smoking-induced transcriptional heterogeneity in the human bronchial epithelium at single-cell resolution. Science Advances 5, eaaw3413.

22. Qiu, F., Liang, C.-L., Liu, H., Zeng, Y.-Q., Hou, S., Huang, S., Lai, X., and Dai, Z. (2016). Impacts of cigarette smoking on immune responsiveness: Up and down or upside down? Oncotarget 8, 268–284.

23. Teixeira, V. H., Nadarajan, P., Graham, T. A., Pipinikas, C. P., Brown, J. M., Falzon, M., Nye, E., Poulsom, R., Lawrence, D., Wright, N. A., et al. (2013). Stochastic homeostasis in human airway epithelium is achieved by neutral competition of basal cell progenitors. eLife 2, e00966.

24. Clayton, E., Doupé, D. P., Klein, A. M., Winton, D. J., Simons, B. D., and Jones, P. H. (2007). A single type of progenitor cell maintains normal epidermis. Nature 446, 185– 189.

25. Watson, J. K., Rulands, S., Wilkinson, A. C., Wuidart, A., Ousset, M., Van Keymeulen, A., Göttgens, B., Blanpain, C., Simons, B. D., and Rawlins, E. L. (2015). Clonal Dynamics Reveal Two Distinct Populations of Basal Cells in Slow-Turnover Airway Epithelium. Cell Reports 12, 90–101.

26. Doupé, D. P., Alcolea, M. P., Roshan, A., Zhang, G., Klein, A. M., Simons, B. D., and Jones, P. H. (2012). A Single Progenitor Population Switches Behavior to Maintain and Repair Esophageal Epithelium. Science 337, 1091–1093.

27. Piedrafita, G., Kostiou, V., Wabik, A., Colom, B., Fernandez-Antoran, D., Herms, A., Murai, K., Hall, B. A., and Jones, P. H. (2020). A single-progenitor model as the unifying paradigm of epidermal and esophageal epithelial maintenance in mice. Nature Communications 11, 1429.

28. Metzcar, J., Jutzeler, C. R., Macklin, P., Köhn-Luque, A., and Brüningk, S. C. (2024). A review of mechanistic learning in mathematical oncology. English. Frontiers in Immunology 15.

29. Procopio, A., Cesarelli, G., Donisi, L., Merola, A., Amato, F., and Cosentino, C. (2023). Combined mechanistic modeling and machine-learning approaches in systems biology – A systematic literature review. Computer Methods and Programs in Biomedicine 240, 107681.

30. Vásquez-Venegas, C., Wu, C., Sundar, S., Prôa, R., Beloy, F. J., Medina, J. R., McNichol, M., Parvataneni, K., Kurtzman, N., Mirshawka, F., et al. (2024). Detecting and Mitigating the Clever Hans Effect in Medical Imaging: A Scoping Review. Journal of Imaging Informatics in Medicine.

31. Díaz-Gay, M., Vangara, R., Barnes, M., Wang, X., Islam, S. M. A., Vermes, I., Duke, S., Narasimman, N. B., Yang, T., Jiang, Z., et al. (2023). Assigning mutational signatures to individual samples and individual somatic mutations with SigProfilerAssignment. Bioinformatics 39, btad756.

32. Tate, J. G., Bamford, S., Jubb, H. C., Sondka, Z., Beare, D. M., Bindal, N., Boutselakis, H., Cole, C. G., Creatore, C., Dawson, E., et al. (2019). COSMIC: the Catalogue Of Somatic Mutations In Cancer. Nucleic Acids Research 47, D941–D947.

33. Alexandrov, L. B., Kim, J., Haradhvala, N. J., Huang, M. N., Tian Ng, A. W., Wu, Y., Boot, A., Covington, K. R., Gordenin, D. A., Bergstrom, E. N., et al. (2020). The repertoire of mutational signatures in human cancer. Nature 578, 94–101.

34. Frankell, A. M., Dietzen, M., Al Bakir, M., Lim, E. L., Karasaki, T., Ward, S., Veeriah, S., Colliver, E., Huebner, A., Bunkum, A., et al. (2023). The evolution of lung cancer and impact of subclonal selection in TRACERx. Nature 616, 525–533.

35. Singh, V. K., Rastogi, A., Hu, X., Wang, Y., and De, S. (2020). Mutational signature SBS8 predominantly arises due to late replication errors in cancer. Communications Biology 3, 1–10.

36. Alexandrov, L. B., Nik-Zainal, S., Wedge, D. C., Aparicio, S. A. J. R., Behjati, S., Biankin, A. V., Bignell, G. R., Bolli, N., Borg, A., Børresen-Dale, A.-L., et al. (2013). Signatures of mutational processes in human cancer. Nature 500, 415–421.

37. Chen, J.-M., Férec, C., and Cooper, D. N. (2013). Patterns and Mutational Signatures of Tandem Base Substitutions Causing Human Inherited Disease. Human Mutation 34, 1119– 1130.

38. Silverman, B. W. (1981). Using Kernel Density Estimates to Investigate Multimodality. Journal of the Royal Statistical Society. Series B (Methodological) 43, 97–99.

39. Hall, P. and York, M. (2001). On the Calibration of Silverman’s Test for Multimodality. Statistica Sinica 11, 515–536.

40. Fisher, N. I. and Marron, J. S. (2001). Mode testing via the excess mass estimate. Biometrika 88, 499–517.

41. Hartigan, J. A. and Hartigan, P. M. (1985). The Dip Test of Unimodality. The Annals of Statistics 13, 70–84.

42. Cheng, M.-Y. and Hall, P. (1998). Calibrating the Excess Mass and Dip Tests of Modality. Journal of the Royal Statistical Society. Series B (Statistical Methodology) 60, 579–589.

43. Ameijeiras-Alonso, J., Crujeiras, R. M., and Rodriguez-Casal, A. (2021). multimode: An R Package for Mode Assessment. Journal of Statistical Software 97, 1–32.

44. Pedregosa, F., Varoquaux, G., Gramfort, A., Michel, V., Thirion, B., Grisel, O., Blondel, M., Prettenhofer, P., Weiss, R., Dubourg, V., et al. (2011). Scikit-learn: Machine Learning in Python. Journal of Machine Learning Research 12, 2825–2830.

45. Lemant, J., Le Sueur, C., Manojlovic, V., and Noble, R. (2022). Robust, Universal Tree Balance Indices. Systematic Biology 71, 1210–1224.

46. Gillespie, D. T. (1976). A general method for numerically simulating the stochastic time evolution of coupled chemical reactions. Journal of Computational Physics 22, 403–434.

47. Alexandrov, L. B. and Stratton, M. R. (2014). Mutational signatures: the patterns of somatic mutations hidden in cancer genomes. Current Opinion in Genetics & Development 24, 52– 60.

48. Toni, T. and Stumpf, M. P. H. (2010). Simulation-based model selection for dynamical systems in systems and population biology. Bioinformatics 26, 104–110.

49. Bernton, E., Jacob, P. E., Gerber, M., and Robert, C. P. (2019). Approximate Bayesian computation with the Wasserstein distance. Journal of the Royal Statistical Society Series B: Statistical Methodology 81, 235–269.

50. Kullbach, S. (1959). Information theory and statistics.

51. Virtanen, P., Gommers, R., Oliphant, T. E., Haberland, M., Reddy, T., Cournapeau, D., Burovski, E., Peterson, P., Weckesser, W., Bright, J., et al. (2020). SciPy 1.0: fundamental algorithms for scientific computing in Python. Nature Methods 17, 261–272.

52. Shannon, C. E. (1948). A mathematical theory of communication. The Bell System Technical Journal 27, 379–423.

53. Nembrini, S., König, I. R., and Wright, M. N. (2018). The revival of the Gini importance? Bioinformatics 34, 3711–3718.

54. Williams, M. J., Zapata, L., Werner, B., Barnes, C. P., Sottoriva, A., and Graham, T. A. (2020). Measuring the distribution of fitness effects in somatic evolution by combining clonal dynamics with dN/dS ratios. eLife 9, e48714.

55. Alexandrov, L. B., Ju, Y. S., Haase, K., Van Loo, P., Martincorena, I., Nik-Zainal, S., Totoki, Y., Fujimoto, A., Nakagawa, H., Shibata, T., et al. (2016). Mutational signatures associated with tobacco smoking in human cancer. Science 354, 618–622.

56. Tomkova, M., Tomek, J., Kriaucionis, S., and Schuster-Böckler, B. (2018). Mutational signature distribution varies with DNA replication timing and strand asymmetry. Genome Biology 19, 129.

57. Werner, B., Case, J., Williams, M. J., Chkhaidze, K., Temko, D., Fernández-Mateos, J., Cresswell, G. D., Nichol, D., Cross, W., Spiteri, I., et al. (2020). Measuring single cell divisions in human tissues from multi-region sequencing data. Nature Communications 11, 1035.

58. Mesa, K. R., Kawaguchi, K., Cockburn, K., Gonzalez, D., Boucher, J., Xin, T., Klein, A. M., and Greco, V. (2018). Homeostatic Epidermal Stem Cell Self-Renewal Is Driven by Local Differentiation. Cell Stem Cell 23, 677–686.e4.

59. Moran, P. A. P. (1958). Random processes in genetics. Mathematical Proceedings of the Cambridge Philosophical Society 54, 60–71.

60. Cole, B. B., Smith, R. W., Jenkins, K. M., Graham, B. B., Reynolds, P. R., and Reynolds, S. D. (2010). Tracheal Basal Cells. The American Journal of Pathology 177, 362–376.

61. Breatnach, E., Abbott, G. C., and Fraser, R. G. (1984). Dimensions of the normal human trachea. American Journal of Roentgenology 142, 903–906.

62. Blokzijl, F., Ligt, J. de, Jager, M., Sasselli, V., Roerink, S., Sasaki, N., Huch, M., Boymans, S., Kuijk, E., Prins, P., et al. (2016). Tissue-specific mutation accumulation in human adult stem cells during life. Nature 538, 260–264.

63. Maughan, E. F., Hynds, R. E., Pennycuick, A., Nigro, E., Gowers, K. H. C., Denais, C., Gómez-López, S., Lazarus, K. A., Orr, J. C., Pearce, D. R., et al. (2022). Cell-intrinsic differences between human airway epithelial cells from children and adults. iScience 25, 105409.

64. Gowers, K. H. C., Hynds, R. E., Thakrar, R. M., Carroll, B., Birchall, M. A., and Janes, S. M. (2018). Optimized isolation and expansion of human airway epithelial basal cells from endobronchial biopsy samples. Journal of Tissue Engineering and Regenerative Medicine 12, e313–e317.

65. Boers, J. E., Ambergen, A. W., and Thunnissen, F. B. J. M. (1998). Number and Proliferation of Basal and Parabasal Cells in Normal Human Airway Epithelium. American Journal of Respiratory and Critical Care Medicine 157, 2000–2006.

66. Chkhaidze, K., Heide, T., Werner, B., Williams, M. J., Huang, W., Caravagna, G., Graham, T. A., and Sottoriva, A. (2019). Spatially constrained tumour growth affects the patterns of clonal selection and neutral drift in cancer genomic data. PLOS Computational Biology 15, e1007243.

67. Sondka, Z., Bamford, S., Cole, C. G., Ward, S. A., Dunham, I., and Forbes, S. A. (2018). The COSMIC Cancer Gene Census: describing genetic dysfunction across all human cancers. Nature Reviews Cancer 18, 696–705.

68. Harrison, P. W., Amode, M. R., Austine-Orimoloye, O., Azov, A. G., Barba, M., Barnes, I., Becker, A., Bennett, R., Berry, A., Bhai, J., et al. (2024). Ensembl 2024. Nucleic Acids Research 52, D891–D899.

69. Piovesan, A., Caracausi, M., Antonaros, F., Pelleri, M. C., and Vitale, L. (2016). GeneBase 1.1: a tool to summarize data from NCBI gene datasets and its application to an update of human gene statistics. Database: The Journal of Biological Databases and Curation 2016, baw153.

70. Benson, D. A., Cavanaugh, M., Clark, K., Karsch-Mizrachi, I., Lipman, D. J., Ostell, J., and Sayers, E. W. (2013). GenBank. Nucleic Acids Research 41, D36–D42.

71. Martincorena, I., Raine, K. M., Gerstung, M., Dawson, K. J., Haase, K., Van Loo, P., Davies, H., Stratton, M. R., and Campbell, P. J. (2017). Universal Patterns of Selection in Cancer and Somatic Tissues. Cell 171, 1029–1041.e21.

72. Denker, J., Gardner, W., Graf, H., Henderson, D., Howard, R., Hubbard, W., Jackel, L. D., Baird, H., and Guyon, I. (1988). “Neural Network Recognizer for Hand-Written Zip Code Digits”. Vol. 1.

73. Mead, A. (1992). Review of the Development of Multidimensional Scaling Methods. Journal of the Royal Statistical Society. Series D (The Statistician) 41, 27–39.

74. Kruskal, J. B. (1964). Multidimensional scaling by optimizing goodness of fit to a nonmetric hypothesis. Psychometrika 29, 1–27.

75. Leeuw, J. D. (1977). Applications of Convex Analysis to Multidimensional Scaling. Recent Developments in Statistics, 133–146.

76. Breiman, L. (2001). Random Forests. Machine Learning 45, 5–32.

77. Vapnik, V. N. (1997). “The Support Vector method”, 261–271.

78. Ben-Hur, A., Ong, C. S., Sonnenburg, S., Schölkopf, B., and Rätsch, G. (2008). Support Vector Machines and Kernels for Computational Biology. PLOS Computational Biology 4, e1000173.

79. Ewens, W. J. (1979). Mathematical Population Genetics.

80. Parsons, T. L. and Quince, C. (2007). Fixation in haploid populations exhibiting density dependence I: The non-neutral case. Theoretical Population Biology 72, 121–135.

