## Supplementary Text and Figures for "Recovery of human upper airway epithelium after smoking cessation is driven by a slow-cycling stem cell population and immune surveillance"

### Supplemental Text

#### Modelling framework

The description of airway basal cell homeostasis given by Teixeira et al.[23] provides a baseline model of the homeostatic backdrop on which somatic evolution can occur in human upper airway stem cell populations. For modelling data from bulk culture of stem cells, we omitted differentiating cells as irrelevant to the long-run clonal dynamics. The observations laid out in the main text are highly suggestive of evolutionary effects. A requirement for Darwinian evolution to take place is heritable phenotypic variation between cells, undergoing selection; this phenotypic variation is commonly projected into a single numerical dimension and given the name fitness. Given the maintenance of homeostasis, the standard model used in cancer of fitness that affects cells' division rates is unsuitable in this instance: instead, a rational and commonly used[54] model is to allow fitness to affect the likelihood of differentiation. By altering a cell's fate distribution (Figure 2A), high fitness can lead a cell on average to increase the number of its descendants within the stem cell pool, by reducing the number of its descendants that will differentiate into (for example) ciliated or goblet cells, contributing to the function of the lung. This notion of a limited selfishness on the part of lineages within stem cell populations in histologically normal tissue, prioritising future generations' numbers over lung function, contrasts with the rapid clonal expansion and metabolic reprogramming observed in cancers.

##### Basic model

In this work, we have used a model for lung basal cell homeostasis over the course of an individual's lifetime, based on their smoking history. In the model, cells divide and are lost to differentiation according to the schema of Clayton et al.[24], presented in Figure 2A, with each division stochastically classified as one of the following:

**symmetric division** into two basal cells;

**asymmetric division** into one basal and one differentiating cell, which is treated within the model as removed from the system (only basal cells are of interest as the data for comparison is derived exclusively from cells with proliferative potential);

**symmetric differentiation** into two differentiating cells, both removed from the system.

During periods in which the modelled individual was smoking, we model an increase in the division rate (lifetime cell turnover is higher in ever-smokers[23]) and an increase in the mutation rate (due to the numerous mutagenic chemicals in tobacco smoke). As such, cells divide at smoking-dependent constant average rate $\lambda_S$ or $\lambda_{NS}$ divisions per year, and acquire mutations on division at rate $\mu_S$ or $\mu_{NS}$ mutations per year. Each division brings with it a random number $M \sim \text{Poisson}\left(\frac{\mu}{\lambda}\right)$ of mutations. In reality, mutations also occur over time independently of cell division, but as tobacco-induced mutations occur predominantly during divisions[55,56] the assumption is made that any other mutations occurring outside of divisions will cause similar dynamics to a commensurate increase in the per-division rate of mutations in the absence of smoke. This assumption has been made in previous work modelling acquisition of mutations in human tissue[57].

In this model, each mutation brings with it an associated fitness change, which can be positive (a somatic-tissue driver mutation, abbreviated here to a "driver mutation" with the caveat of its philosophical difference from a cancer driver mutation noted), negative (a deleterious mutation) or null. The distribution of fitness effects is laid out below (section "Distribution of fitness effects summary"). A cell's fitness at any given point in time is the sum of the fitness changes associated with each mutation it has inherited or acquired. This value is additively normalised over the entire population of cells, to represent that this is a competitive advantage in cell fate imbalance rather than a purely cell-intrinsic decision – this represents the continued rule of the organism over the cells; they can only compete with their neighbours to gain selective advantage by reducing their share of a maintained level of differentiation. A constant population of cells is maintained (by different methods in the well-mixed and spatial implementations).

A cell's fitness value is incorporated into the cell fate distribution on division by the rescaled logistic function

$$S_r : \mathrm{Reals} \to [-r, r], \quad f \mapsto r \times \frac{1 - e^{-f}}{1 + e^{-f}},$$

where $r < 0.5$ is the symmetric differentiation probability. The projected value is added to the probability of symmetric division, and subtracted from the probability of symmetric differentiation (see below). This induces a "tilt" in division (schematised in Figure 2A) towards proliferation within the basal cell population for cells with positive fitness, or towards differentiation out of the basal cell population for cells with negative fitness. Divisions occur at constant rate, and have cell fate distribution as follows (B representing a basal cell with fitness f, D cells on a terminal differentiation pathway):

$$B \to \begin{cases} B + B \text{ with probability } r + S_r(f)) \\ B + D \text{ with probability } 1 - 2r \\ D + D \text{ with probability } r - S_r(f)) \end{cases}$$

The number of mutations occurring on each cell division is modelled by the Poisson distribution, as an approximation to the binomial distribution over all positions in the genome. As the rate of mutations increases during smoking, smoking signatures can be simulated by subdividing the mutation count on any division $M \sim \mathrm{Poisson}\left(\frac{\mu}{\lambda}\right)$, into $M = M_{NS} + M_S$, where

$$M_{NS} \sim \mathrm{Poisson}\left(\frac{\mu_{NS}}{\lambda_{NS}}\right),$$

$$M_S \sim \begin{cases} \mathrm{Poisson}\left(\max\left(\frac{\mu_S}{\lambda_S} - \frac{\mu_{NS}}{\lambda_{NS}}, 0\right)\right) & \text{if smoking} \\ 0 & \text{otherwise.} \end{cases}$$

The mutation rate per division $\frac{\mu}{\lambda}$ should be higher during smoking; in the edge case of it being smaller (i.e. if the division rate increases more than the division rate during smoking, a biological implausibility given the existence of smoke-associated mutational signatures), no smoking signature mutations would be observed and hence $M_S = 0$.

##### Quiescent subpopulation hypothesis

The hypothesised subpopulation of cells which divide more slowly is modelled as an additional compartment of a separate cell type, also included in the stem cell pool. These have the same mutation rate per division, but a lower division rate than the main cell population (by some multiplicative factor between 0 and 1, a parameter of the model), increased by the same factor during smoking. On division, they produce one quiescent cell and one basal cell, so as to provide a constant supply of less-mutated cells into the main population. These “cells” correspond either to individual slow-dividing basal cells or to some larger biological entity such as a small gland, in which the time to fixation for any new mutation is low, and there is a low overall division rate (see section “Quiescent gland fixation probability”).

An additional “sub-hypothesis” is included that the quiescent cells are themselves protected from smoke effect, and therefore that their division and mutation rates are not augmented as much as that of other basal cells.

##### Protected subpopulation hypothesis

The hypothesised protected sub-population is also modelled as an additional cell compartment, this one comprising cells which divide in a manner identical to the main population but which are less affected by smoking. These could represent a physically protected niche of cells in which basal cells behave identically to outside of the niche but without as strong an effect from smoking.

This reduction in the effect of smoking is modelled linearly: if normal cells divide at rate $\lambda_{\mathrm{NS}}$ and mutate at rate $\mu_{\mathrm{NS}}$, and during smoking these change to $\lambda_{\mathrm{S}} > \lambda_{\mathrm{NS}}$ and $\mu_{\mathrm{S}} > \mu_{\mathrm{NS}}$, then with protection coefficient $\alpha$, a protected cell would have equivalent rates

$$\lambda_{\mathrm{NS}}^{(\alpha)} = \lambda_{\mathrm{NS}} \qquad \lambda_{\mathrm{S}}^{(\alpha)} = \lambda_{\mathrm{NS}} + \alpha(\lambda_{\mathrm{S}} - \lambda_{\mathrm{NS}})$$
$$\mu_{\mathrm{NS}}^{(\alpha)} = \mu_{\mathrm{NS}} \qquad \mu_{\mathrm{S}}^{(\alpha)} = \mu_{\mathrm{NS}} + \alpha(\mu_{\mathrm{S}} - \mu_{\mathrm{NS}}).$$

This same linear reduction in effect is applied to mutation rate and division rate, but not rate of immune killing as this would not be affected by exposure to the airway.

##### Differential immune response hypothesis

Immune response to somatic mutations is already partially factored into the model implicitly via the distribution of mutational fitness effects: most simulated mutations are deleterious to varying degrees, which may be partially attributed to the action of the immune system. However, in order to explain the observed recovery of the near-normally mutated sub-population of cells after the cessation of smoking (shown in Figure S1), a mechanism is required to represent the immune predation of highly mutated cells with reduced efficacy during smoking. This is implemented as a constant stochastic death rate for each cell, proportional to the number of mutations it has acquired, and reduced by some coefficient (a parameter) during smoking.

##### Smoking-addicted driver mutation hypothesis

To model the hypothetically increased efficacy of driver mutations during smoking, a linear augmentation is applied to the fitness effect of all driver mutations during smoking, multiplying each by some global coefficient $1 + \gamma$, $\gamma > 0$. Smoking has a broad effect on the lung environment[21], and due to difficulties in estimating the fitness effects of somatic mutations there is a lack of understanding of the precise effects this change in environment has on the distribution of fitness effects. Motivated by the observed rebalancing of populations by mutational burden after stopping smoking[3], this hypothesis posits that the change may be to add to the cell fate bias-inducing mutations causing increased proliferation in the stem cell compartment.

##### Distribution of fitness effects summary

In order to randomly allocate fitness values to new mutations, a distribution is required to approximate that of fitness effects in somatic evolution *in vivo*. While the distribution of fitness effects (DFE) in somatic evolution in human epithelia is largely unmapped, first forays into parsing it from genomic sequencing of healthy skin and oesophagus[8,9] show a landscape of mostly neutral mutations, with a small tail of driver mutations that confer a selective advantage[54]. Previous analyses have demonstrated that the exact shape of DFE often has no bearing on the outcome of simulation analyses[54].

In this study, we define the DFE as follows:

- Most mutations have no effect on a cell's fitness. The fraction of mutations that have any effect is taken to be 10% (see below section "Distribution of fitness effects").
- Of those that do have a fitness effect, the probability of the effect being positive is 19.6% for spatial simulations, and 0.78% for non-spatial simulations, in order to align the rate of significantly fitness-affecting driver mutations with the rate of cancer driver mutations in the genome (see below section "Distribution of fitness effects").
- The absolute value of the fitness effect is drawn from an exponential distribution (as used previously[54]), whose scale is a parameter of the model and is set as either 0, 0.05 or 0.1 to allow for fitting the presence and degree of selection to the data (values selected via exploratory simulations; see below section "Distribution of fitness effects").

Cells accumulate multiple fitness-affecting mutations over the course of a human lifetime. The fitness of a cell is taken to be the sum of its mutations' fitness values, which is additively normalised and projected via a sigmoid function onto the relevant range $[-r, r]$, as defined above.

##### Integration into spatial lattice

Human airway basal cells maintain constant contact with the basement membrane, so they can be modelled in two dimensions (Figure 2B). In order to maintain a constant density of basal cells across the epithelium, it has been shown in the murine epidermis via longitudinal imaging that stem cells divide in response to a nearby stem cells' differentiation[58], over a time scale of around a day. A parsimonious way to model a spatial system of cells with such a homeostatic mechanism is via a Moran-like model[59], with a two-dimensional lattice in which each point represents a single basal cell, and each symmetric differentiation or cell death is immediately followed by a symmetric division by an adjacent cell, chosen stochastically with weighting by division rate (and hence by cell fitness), to fill the gap.

We implemented this augmented model via a Gillespie algorithm[46] in which the symmetric differentiations of all cells, as well as their asymmetric divisions, occur at constant rates. This assumes that the time between divisions for each cell is exponentially distributed; in reality there may be internal and external factors that lead to cell divisions with a different timescale. Each symmetric differentiation or immune cell death then leads to a selection from the 4 immediate neighbours of that cell in the square lattice, weighted by their symmetric division rates as set out above, to divide symmetrically to fill the gap. In this way, each division type will occur on average at the rates set out in the homeostatic model[23], and higher fitness clones will have a higher chance of expanding into gaps created by loss of cells.

We integrated each modelling hypothesis into the simulation structure, taking into account its biological motivation:

**Quiescent subpopulation** Quiescent cells are modelled as residents of genetically homogeneous glands, which form a sparser lattice below the basal cell lattice. If the basal cell immediately above a quiescent gland is lost, then the quiescent gland can compete to produce a new basal cell daughter cell to fill the gap. Divisions of cells within glands without escaping is modelled as a constant rate of stochastic mutation accumulation, another of the events occurring via the Gillespie algorithm, with a fixation probability selection filter applied to mutations occurring outside of basal daughter cell production (see section "Quiescent gland fixation probability"). The number of cells contained within each gland is a parameter of the model; setting this to 1 models single cells rather than glands.

**Protected subpopulation** The subpopulation of cells less affected by tobacco is modelled by diamond patches of lattice (equivalent to circles within the canonical metric space on the lattice) in which smoking has reduced effect on cell turnover and mutation rate. While no migration is included in the model, daughter cells of protected cells may be unprotected, and vice versa. The size of these regions can be varied, but in order to reduce the dimensionality of the parameter space was fixed at a radius of 7 cells, leading each region to cover 113 cells.

**Differential immune response** This hypothesis is not affected by the transition to a spatial system. Cell deaths due to the immune system are another event modelled with the Gillespie algorithm. It is feasible to suppose that immune regulation would in fact be triggered only by a sufficiently large clone; the effect of this is not investigated here for model parsimony.

**Smoking driver augmentation** This hypothesis is also uncomplicated by implicit spatial assumptions, and can be transplanted unchanged into the spatial implementation.

There are approximately 5000 basal cells around a cross-section of an average human trachea[60,61]. In order to model the true cylindrical topology on which these somatic evolutionary processes take place, 25 million cells would therefore need to be simulated, which is computationally intractable. Instead, a small subsection of the epithelium is modelled; in this context the local topology of a cylinder is that of a flat plane, which is modelled by a torus to remove edge effects.

This implementation was written in the Julia programming language, version 1.9.0. Mutations and divisions were stored in bespoke data structures such that phylogenies could be reconstructed, considering only lineages visible from extant or subsampled cells.

##### Simulation parameter scales

In order to make inference from data in the light of simulations with unknown parameter values, meaningful prior distributions are required for the model and for each parameter, derived from literature if possible or uninformative otherwise. An uninformative uniform prior distribution over models is taken, and the parameters' prior distributions are given in Table S3 and explained in the remainder of this section.

Parameters naturally vary on different scales in biological systems. Continuously-varying parameters are given normally distributed priors on their relevant varying scale. The given mean and standard deviation are in the "true" scale on which the parameters are used; an appropriate normal distribution in the varying scale is calculated such that the requisite mean and standard deviation are achieved. In the case of a log-scale, this has an analytical solution: for true-scale mean $\mu$ and standard deviation $\sigma$, log-scale mean and standard deviation must be

$$\mu^* = \log(\mu) - \frac{1}{2}\log\left(1 + \frac{\sigma^2}{\mu^2}\right) \tag{3}$$

$$\sigma^* = \sqrt{\log\left(1 + \frac{\sigma^2}{\mu^2}\right)}. \tag{4}$$

In the case of a logit varying scale, in which parameters vary on a scale accessed by the function

$$\text{logit} : \text{Reals} \to [0, 1], x \mapsto \log\left(\frac{x}{1 + x}\right),$$

there is no such closed-form solution for $\mu^*, \sigma^*$. As such, we used the *fsolve* function from the Python package *scipy.optimize* (version 1.12.0,[51]) to numerically solve for appropriate values, with the mean and standard deviation of a random variable $Y \sim \text{Normal}(\mu^*, \sigma^{*2})$ after *expit*-transformation calculated using Monte Carlo approximation

$$E\left[\text{expit}(Y)^n\right] \approx \frac{1}{K-1}\sum_{i=1}^{K-1} \text{expit}(\Phi^{-1}_{\mu^*,\sigma^*}(i/K))^n \quad \text{for large } K$$

for $n = 1, 2$, where $\Phi_{\mu^*,\sigma^*}$ is the cumulative distribution function of Y and E represents expectation. K is augmented until convergence.

##### Simulation parameters and prior distributions

The prior mean values for mutation rates are based on estimates from the same data by Yoshida, Gowers et al[3], which are broadly in line with previous estimates for other tissues[62].

The coefficient of the distribution of fitness changes is given a discrete prior in order to allow for non-zero probability of being exactly 0 to signify no selection: it takes values 0, 0.05 and 0.1 with probabilities $\frac{1}{2}, \frac{1}{4}, \frac{1}{4}$ respectively. This provides an uninformative prior on whether selection exists, and a separate uninformative prior on the scale of fitness changes. The choice of these values was made by running small simulations with different values; this analysis is shown in the below section “Distribution of fitness effects”.

Values for division rates while smoking and not smoking are estimated from published work and given narrow priors around these values. Teixeira et al[23] found that the cell turnover rate $2r\lambda$, where r is the symmetric division probability and $\lambda$ is the total division rate (over all division types), has values:

$$2r\lambda = \begin{cases} 1.0 \pm 0.5 \text{ /year while not smoking} \\ 1.8 \pm 0.4 \text{ /year while smoking.} \end{cases}$$

The assumption is made that the value r is unaffected by smoking, and instead the increase in turnover during smoking is brought about entirely by an increase in the division rate $\lambda$. As such, the division rate augmentation due to smoking is given mean value $\frac{1.8}{1.0} - 1 = 0.8$.

Watson et al.[25] used lineage tracing experiments in mouse models to demonstrate, using the same basal cell homeostasis model as Teixeira et al., a division rate of one every 11 days ($\pm$4). This translates to 365.25/11 = 33 (to two significant figures, $\pm$ 15) divisions per year. While this is a readout of mouse characteristics rather than human, no estimates were found for *in vivo* human lung basal cell division rate and *in vitro* estimates of 166 divisions per year (doubling every 2.28 days)[63] should be considered an upper bound in response to damage rather than a base level.

Given that the hypothesised quiescent subpopulation has not been identified, and would only serve the purpose of replenishing the main “workhorse” basal cell population over time, it is not likely to form a large proportion of cells. As such, the fraction of cells which are quiescent has a low prior mean of 10%, but a broad standard deviation of 8%. The number of divisions per year for each quiescent cell has a broad log-normal prior centred around 4, reflecting an unknown but significantly reduced division rate, and the ambient quiescent mutation rate modelling within-gland divisions has a broad prior around half the rate of non-quiescent cells, ie 16.5 divisions per year, so as to remain significantly less mutated than the main basal cell population.

Uninformative priors are used for the fraction of cells which are protected from the effects of smoking, the coefficient by which they are protected, the coefficient by which quiescent cells are protected in addition to dividing slower, and the coefficient by which the action of the immune system decreases during smoking.

The rate at which the immune system kills cells, measured in deaths per mutation per year, is challenging to estimate directly from existing data. However, by running simulations with default values for other parameters, the rate of cell turnover from this immune mechanism can be compared with the level of turnover previously determined in the human lung epithelium[23], 1.0$\pm$0.5 per cell per year. Taking $\frac{1}{4}$ as a reasonable prior mean fraction of this turnover, the value of 0.00025 was selected (see Figure Figure 2G for the immune death rate at this selected value, with other parameters at prior mean values). Given the loose link to experimental findings, this is given a broad log-normal prior distribution.

The coefficient by which fitness is augmented during smoking (within the smoking driver hypothesis) is difficult to estimate *a priori*. One can go some way to estimating its scale by considering the chosen distribution of fitness effects (see section “Distribution of fitness effects”): this distribution will naturally lead to around four-fold more deleterious mutations than advantageous mutations. A driver fitness augmentation above 3 would lead to the average mutation having an advantageous effect, which seems to be evidenced by the more-mutated bulk of cells in current-smokers. As such, a broad prior around 3 is set for this parameter.

#### Sampling from simulations

When comparing a simulation with the observed scWGS dataset, a subsample of surviving cells at the end of the simulation (representing the time of the biopsy being taken) is required to represent the process by which a piece of tissue in the lung is sub-selected down to only at most 60 sequenced cells. The three broad stages of sub-selection in the sampling process for the Yoshida et al. study (from which most inference is drawn, as they sequenced more cells for each patient) are as follows:

- The biopsy being taken from one particular area of lung epithelium
- Only certain cells successfully being cultured
- Only certain bulk cultures being sent for sequencing.

The first of these is a limitation of this work: by default, for computational reasons, these simulations model only 1600 basal cells from each patient's epithelium. In reality, the kind of biopsy samples used by Yoshida et al. contain between 50,000 and 100,000 cells[64], of which an average of 11.6% expressed EPCAM and were thus classified as epithelial cells. 30% of this number (the proportion of basal cells in the upper airway epithelium[65]) takes the figure to between 1740 and 3480 basal cells per biopsy. Simulations of 10,000 cells demonstrate minimal differences from simulations of 1600 cells (see below), providing reassurance that this limitation is minor in scope and unlikely to change conclusions.

The second subselection stage, that of culturing, is likely to be insubstantial as a bottleneck: 15-40% of each patient's cells cultured successfully to a level where they could have been sequenced. This aligns with the estimate of approximately 30% of lung epithelial cells being basal cells[65], suggesting a high colony forming proportion among the basal cells (given that cells on a differentiation pathway would be unlikely to successfully culture).

The third subselection stage defines the subsampling schema used in this work: a specified number (generally around 50) of single-cell-derived cultures were prepared for WGS, selected entirely at random from those that had successfully cultured. We therefore draw cells at random (without replacement) from those alive at the time of sampling, until the number of cells sequenced is reached. This subsample can be used to construct mutational burden distributions and phylogenies as required. A spatial subsampling procedure, as has been performed in previous simulation studies[66], would misinterpret the data sampling procedure: while of course the biopsy represents a spatial subsection of the lung, the entire simulation models an area of tissue smaller than the biopsy. Thus, the only relevant subsampling occurs after sampling and is spatially agnostic.

#### Hypothetical paradigms

On top of the basic model of somatic evolution in lung homeostasis, the hypotheses described in the Introduction are modelled as "modules" that can be independently activated, in order to investigate the impact of their activation, individually or in combination, on the ability of simulations to recapitulate the true data. The subset of hypotheses that are activated represents the full paradigm of lung homeostasis in consideration: as such, in this work we refer to such subsets as "hypothetical paradigms" or "paradigms" for brevity.

A trade-off arises here between simplicity and completeness: these hypotheses interact in interesting ways, and it may well be that some combination of them is the best paradigm to explain the dynamics described in the previous section and summarised in this one, but inference procedures and manual inspection require some degree of parsimony to understand what each hypothesis is contributing. As such, analyses and plots are made using one of two sets of hypothetical paradigms: all 24 combinations, or the "protection-selection" set of the 9 paradigms in which at most one hypothesis from each of the two pairs is active. To clarify, a paradigm (subset of the four hypotheses activated) is included in the protection-selection set of paradigms if and only if it contains at most one hypothesis attempting to explain the existence of near-normally mutated cells in lifelong smokers (quiescent or protected sub-population), and at most one hypothesis attempting to explain the recovery in the lungs of ex-smokers (differential immune response or smoking-addicted drivers). Note that the use of "protection" here is separate from the protected paradigm: here, "protection" means we need some way to explain why some cells were not affected, while the "protected" hypothesis is a particular explanation for this.

The names, abbreviations and brief summaries of the hypotheses are given in Table S2.

#### Non-spatial underdispersion

This section describes an initial attempt to model the basal stem cell population in a well-mixed manner. This model did not recapitulate the dynamics adequately, due to rapid clonal expansion leading to under-dispersion in mutational burden distributions, and is described here for completeness.

The theoretical model described in Methods was initially implemented in a discrete-time agent-based framework, similarly run over the course of each patient's smoking record.

Within this implementation, each year is divided into s simulated time steps (365 by default), and each cell divides, on average, $\lambda_S$ times per year of smoking and $\lambda_{NS}$ times per year not smoking. Similarly, $\mu_S$ and $\mu_{NS}$ represent the mutation rates within each cell per simulated year. These rates are achieved on average by a constant division rate per step and mutation rate per division, with each cell dividing a random number $D \sim \text{Poisson}\left(\frac{\lambda}{s}\right)$ times at each time-step.

Cells compete in a non-spatial manner, with compartments (standard, protected, quiescent, etc.) held separate from each other and competition allowed between all cells within each compartment. Divisions in the quiescent compartment feed cells into the standard compartment; if the quiescent and protected hypotheses are both active, then the protected compartment has a separate quiescent compartment of its own, also protected from smoking effect by the same factor and feeding daughters cells exclusively into the protected compartment.

Homeostatic consistency of population size is implemented differently to the spatial simulation system described in Methods, with a carrying capacity N enforced via cell fitness normalisation: rather than normalising to 0 fitness, which corresponds to equal probability of symmetric division or symmetric differentiation (see Figure 2A), cell fitnesses are normalised to have mean

$$\delta \log\left(\frac{N}{n}\right),$$

where $\delta$ is a hyperparameter controlling the strength of this dynamic normalisation and n is the current number of basal cells. This pushes the cell population to revert towards the original population size, by dividing or differentiating more frequently. Biologically, this represents cell signalling to increase or decrease the basal cell population rather than crowding (as the majority of cells in the pseudostratified epithelium of the upper airway are not basal cells).

Figure S3A shows significantly narrower mutational burden distributions than was observed in Figure S1C., Omitting cell-cell interactions and considering only mutational burden accumulation (see Theoretical dispersion below), we find a metric of dispersion relevant to the model; this metric is significantly lower in well-mixed simulations than theoretical predictions without cell-cell interactions (Figure S3B)), and than in the true data (Figure S3C)).

**Theoretical dispersion**

Here we provide a simplified version of the simulation model described in Methods which is analytically tractable and can thus be interrogated for expected dispersion.

Ignoring all proliferation, differentiation and death, and considering only the mutation element of these simulations, creates a simpler model in which the distribution of mutations follows a compound Poisson distribution: the number of divisions D accumulated in a particular cell over the course of a simulation is a sum of the Poisson-distributed random variable numbers of divisions at each step, so follows a Poisson distribution itself; write $D \sim \text{Poisson}(a)$ for some $a > 0$. The number of mutations accumulated over the course of those D divisions is then

$$M = \sum_{i=1}^{D} X_i, \text{ where}$$

$X_1, X_2, \ldots \overset{\text{i.i.d.}}{\sim} \text{Poisson}(b)$ for some $b > 0$ are the numbers of mutations on each division.

Indeed, the mutation counts $\{m_1, \ldots, m_n\}$ in the different simulated cells are, within this approximation, n independent, identically distributed (IID) draws of this random variable M. Analysis of the distribution of M can hence provide insight into the mutational burden distribution in the absence of simulated cell-cell interactions.

The mean and variance of M are as follows:

$$\begin{aligned} E[M] &= E\left[E[M|D]\right] \text{ by the law of total probability} \\ &= E[D \times b] \\ &= ab, \end{aligned}$$

$$\begin{aligned} \text{Var}(M) &= E_D\left[\text{Var}_M(M|D)\right] + \text{Var}_D\left(E_M[M|D]\right) \text{ by the law of total variance} \\ &= E_D[bD] + \text{Var}_D(bD) \\ &= ab(1 + b). \end{aligned}$$

As such, this approximation predicts that the standard deviation of mutational burden would be a constant multiple $\sqrt{1+b}$ of the square root of the mean, where b, the mean number of mutations per division, is, for default parameter values,

$$b = \frac{\text{mutations per year}}{\text{divisions per year}}$$
$$= \begin{cases} 25/33 \approx 0.76 \text{ while not smoking} \\ 100/59.4 \approx 1.7 \text{ while smoking,} \end{cases}$$

giving

$$\sqrt{1+b} = \begin{cases} 1.33 \text{ while not smoking, or} \\ 1.64 \text{ while smoking.} \end{cases}$$

This then gives a useful quantity to consider as the "dispersion" of a collection of mutational burden counts over a set of cells in one (simulated or real) lung: the standard deviation as a fraction of the square root of the mean, a value conceptually akin to the coefficient of variation. In a setting with no cell-cell interaction, this would have a value around 1.33 for never smokers and somewhere between 1.33 and 1.64 for ever smokers.

#### Effects of hypotheses on default-parameter-value simulations

Besides the mechanistic learning inference procedure described in the main text, the behaviour of simulations alone can be studied. In particular, the effects of the hypotheses on somatic evolutionary dynamics can be studied in isolation, without direct comparison to the observed scWGS dataset.

A natural consideration in the study of the effect of smoking is the rate of accumulation of mutations. Figure S4A demonstrates that the rates of mutation accumulation broadly align with mutation rates of individual cells in the absence of hypotheses activated (the "base" paradigm), with a slight reduction likely attributable to negative selection (notably this selection occurs in reverse in the presence of the smoking driver hypothesis during smoking). The quiescent hypothesis has a strong effect in reducing the accumulation rate at all smoking history stages, a behaviour that is increased synergistically in combination with the immune response hypothesis.

Fitting Gaussian mixture models to these simulations reveals Figures S4B, S4D a high proportion of simulations assigned a two-component mixture model in all combinations of hypotheses, with diverse profiles of bimodality by smoking history. Smoking histories are also reflected in the tree balance metric $J_1$[45] (Figure S4C).

Together, these results demonstrate that while differing combinations of hypotheses qualitatively appear similar to the true data by certain summaries, they are diverse with respect to other metrics. This highlights the importance of quantitative mechanistic inference, of the form presented in the main text.

#### Effect of larger simulations

To investigate the effect of simulation size on outputs, we ran simulations of 10,000 cells rather than 1600. Due to computational constraints, only one replicate of each simulation was run at this size, with the same default parameter values across all paradigms.

The main concern with the size of simulations was that the dispersion (see below) would be low, as a result of not having a sufficiently large effective sample size of the distribution of possible independent paths a collection of cells could take (including interactions). A sufficiently large simulation will have cells far enough apart that, in practice, they do not interact over the course of the simulation: this therefore provides, in effect, multiple independent samples of the distribution of cells' interactions within a small neighbourhood of cells, increasing the dispersion up to a limit of what the variance would be between distinct simulations. Figures S3F–S3G show the dispersion values of the larger simulations, which are comparable to those shown in Figures S3D–S3E.

#### Distribution of fitness effects

To decide the proportion of fitness-affecting mutations that should have positive (driver) mutation effects, it is noted that since cancer driver genes often overlap with somatic tissue drivers[54], a reasonable estimate for the proportion of mutations carrying a non-negligible positive fitness effect may be taken as the proportion of mutations in the human genome which are identified cancer driver mutations.

The latter may be estimated from the literature: studies of large aggregated datasets of genomic sequencing of cancers have identified 719 cancer driver genes[67] out of a total of approximately $20,000$ protein-coding genes[68], which in turn make up 1.74% of the human genome[69]. By combining the codon frequencies in the human genome[70] with the fraction of single nucleotide substitutions for each codon which are synonymous, an estimate can be made that mutations in the human genome have a 77% chance of being non-synonymous. Further to this, an estimate has been made in breast cancer samples[71] that 49% of non-synonymous mutations in cancer driver genes are positively selected for. Together, these provide a broad estimate for the fraction of mutations to the human genome that are "significant drivers" (using the notation $\tilde{P}(\cdot)$ for an empirical estimate of a probability):

$$\begin{aligned}\tilde{P}(\text{significant driver}) =& \tilde{P}(\text{in coding gene}) \times \tilde{P}(\text{in driver gene} \mid \text{in coding gene}) \times \quad (5a)\\ & \tilde{P}(\text{non-synonymous}) \times \tilde{P}(\text{driver} \mid \text{non-synonymous in driver gene})\\ =& 0.0174 \times \frac{719}{20000} \times 0.77 \times 0.49\\ =& 2.36 \times 10^{-4} \quad (5b)\end{aligned}$$

The following section describes a simplified model which can be used to define a fitness change threshold $x > 0$, such that any mutation with fitness change above this threshold can be labelled as a "significant driver." The fraction of such mutations can thus be fit to the above estimate for the fraction of mutations to the human genome which are known cancer drivers, which specifies a value for the probability $m$ that a random mutation with non-zero fitness effect will have positive fitness effect.

**Continuous-time Markov Chain approximation**

The aim of this model is to assess the effect, within the spatial simulation model, of a mutation of fitness effect f on a clone's subsequent survival probability and growth. This model can then be used to find a threshold for the value f above which a mutation is considered to be a "significant somatic driver", which can then be used to fit the DFE to the rate of known driver mutations in the human genome.

To model this, we envision a lattice in which a single cell is given such a mutation at time $t = 0$, with no other mutations taking place.

First we assume that the clone will grow in a roughly circular shape, and note that the number of lattice grid points adjacent to such a contiguous, near-circular clone of n cells is approximately equal to $2\sqrt{\pi n}$, as is the number of points within the clone that have any directly adjacent external (unmutated) cells. It is also noted that most cells outside the clone will have at most one adjacent cell in the clone, and most cells within the clone will have at least 3 adjacent cells within the clone.

The size of the clone at time $t > 0$ can therefore be approximated by a birth-death process, a continuous-time Markov Chain (CTMC) on the non-negative integers with only adjacent transitions, with 0 as an absorbing state and transition rates

$$\begin{aligned} b_n &= \#\{\text{adjacent cells}\} \times \text{rate of unmutated cell symmetric differentiation} \\ &\quad \times \text{P(mutated cell fills gap)} \\ &\approx 2\sqrt{\pi n} \times r\lambda \times \frac{1 + \sigma(f)}{4 + \sigma(f)} \quad (6a) \\ d_n &= \#\{\text{boundary cells}\} \times \text{rate of mutated cell symmetric differentiation} \\ &\quad \times \text{P(unmutated cell fills gap)} \\ &\approx 2\sqrt{\pi n} \times r\lambda(1 - \sigma(f)) \times \frac{1}{4 + 3\sigma(f)}, \quad (6b) \end{aligned}$$

where $\sigma(f) := \frac{1-e^{-f}}{1+e^{-f}}$ is a sigmoid function projecting fitness onto $(-1, 1)$.

From this approximation, we aim to find two properties of different positive values of f, in order to find a reasonable value for what may be termed a "significant" driver mutation to compare with the rate of known driver mutations:

1. The expected clone size after t years
2. The probability of a clone dying out within t years

As noted by Williams et al[54], the non-spatial (well-mixed) version of this simplified model has rates

$$b_n = \frac{1 + \sigma(f)}{2} \quad (7a)$$

$$d_n = \frac{1 - \sigma(f)}{2}, \quad (7b)$$

and is solvable for the two quantities listed above:

$$\mathrm{E}\left[\text{Clone size at time t}\right] = \frac{(1+\sigma(f))e^{2r\lambda\sigma(f)t} - (1-\sigma(f))}{2\sigma(f)} \tag{8a}$$

$$\mathrm{P}\left(\text{Extinction in t years}\right) = \frac{(1-\sigma(f))(e^{2r\lambda\sigma(f)t} - 1)}{(1+\sigma(f))e^{2r\lambda\sigma(f)t} - (1-\sigma(f))} \tag{8b}$$

This simplified system, in both its spatial and non-spatial forms, can be investigated using Gillespie simulations[46]. Running 100,000 simulations for a range of values of $\sigma(f) \in [0, 1]$, the projected fitness value allows estimation of bootstrap confidence intervals for the values of interest (expected clone size and extinction probability, defined above). The mean values match the analytical solution above in the non-spatial setting (Figures S5A–S5B).

We can use these bootstrap samples to specify a threshold for $\sigma(f)$ in each setting above which a mutation is a "significant" driver. This is taken as the level of $\sigma(f)$ at which the mean number of cells after 60 years is above the 95$^{th}$ percentile of the equivalent value in the neutral setting $F = 0$. We note that at the threshold values of $\sigma(f)$, the probability of a new clone surviving the simulated 60 years is non-negligible at around 7% for the non-spatial model and 26% for the spatial model (Figures S5C–S5D).

##### Inference of fitness change distribution parameter

Finally, we use these thresholds for $\sigma(f)$ to infer values for the probability m of a fitness-altering mutation bringing a positive fitness change, by comparing with the estimate $\tilde{P}$(significant driver) derived above for the probability that a random mutation in the human genome will be a cancer driver mutation. We note that, within the fitness change distribution as laid out in Methods,

$$\mathrm{P}(\text{significant driver}) = p \times m \times \mathrm{P}(\text{significant driver}|\text{positive fitness change}),$$

where $p = \mathrm{P}(\text{non-zero fitness change})$ and $m = \mathrm{P}(\text{fitness} > 0|\text{non-zero fitness change})$ are parameters. Rearranging and substituting the estimate $\tilde{P}$(driver) for its counterpart, we find

$$m = \frac{2.36 \times 10^{-4}}{p \times \mathrm{P}(\text{significant driver}|\text{positive fitness change})},$$

Given that the magnitude of a non-zero fitness change is modelled by the distribution Exp($\alpha$) where $\alpha$ is the fitness change scale (a parameter), we can calculate that for a significance threshold value x for the projected fitness $\sigma(f)$, within the model's fitness change distribution,

$$\mathrm{P}(\text{significant driver}|\text{positive fitness change}) = e^{-\frac{\sigma^{-1}(x)}{\alpha}} = \left(\frac{1-x}{1+x}\right)^{\frac{1}{\alpha}}.$$

This formula is used within the simulation framework to calculate the value of m, the probability of a mutation having positive fitness change given that it has non-zero fitness change, from the fitness change scale $\alpha$ and fitness change probability p.

The remaining two undetermined parameters of the fitness change distribution are the fitness change scale $\alpha$ and the probability of non-zero fitness change p. These parameters have certain constraints, both on their values and on the simulations they produce:

- Fitness change scale should be sufficiently large to have an effect
- Fitness change scale should not be sufficiently large that different populations have deterministic interactions (for example quiescent and non-quiescent populations)
- Mutations should be on average deleterious, as revealed by *ex vivo* competition assays by Maughan et al.[63] revealing a decrease in proliferative fitness of human upper airway stem cells with age.

To employ these constraints, a grid search was performed over values for p and $\alpha$. At each set of values, a simulation was performed using the spatial simulation framework set out in Methods of 1600 cells in the lungs of an 80-year-old never smoker. Values of (p, $\alpha$) for which the mean unnormalised fitness was positive were discarded (Figure S5E). Values p = 0.1, $\alpha$ = 0.05 thus satisfy the third constraint; they also satisfy the first (Figure S5F) and second (Figure 2E). As such, these values were chosen.

The value that this Appendix set out to calculate, the probability within the chosen fitness change distribution that a mutation with non-zero fitness effect should have positive fitness effect, is thus

$$m = \begin{cases} 0.78\% & \text{in non-spatial simulations} \\ 19.6\% & \text{in spatial simulations} \end{cases}$$

Therefore, a full description of the fitness change distribution in spatial simulations is a random variable drawn as follows:

$$F \sim \begin{cases} \text{Exp}(0.05) & \text{with probability } 0.196 * 0.1 \\ -\text{Exp}(0.05) & \text{with probability } (1 - 0.196) \times 0.1 \\ 0 & \text{otherwise.} \end{cases}$$

#### Identifiability analysis

As well as comparing a simulated cohort with the observed cohort to evaluate the simulation paradigm, one can compare simulated cohorts with each other in order to assess the ability of differing metrics and combinations of metrics to distinguish paradigms or parameter value changes from their outputs. This is the essential idea of an identifiability analysis, and it provides us with three useful readouts from the combination of our model, implementation and set of comparison metrics:

- Given an output of a simulation, how confident can we be of the simulation paradigm that generated it (using the metrics defined above)?
- Does this confidence depend on the paradigm that generated it?
- In making this reverse inference (taking a simulated cohort to the putative paradigm that produced it), which metrics are useful?

To assess the metrics described in the previous section against each other, we generated a group of synthetic cohorts, generated in different paradigms and with parameter values drawn randomly from their prior distributions (Table S3). Note that this analysis differs from that presented in Figure 2 and Figures S3–S4 in this regard: rather than taking default parameter values to inspect the general behaviour of simulations, here we drew random parameter values from their prior distributions to investigate the distribution in output space.

Three similarity levels could then be defined between pairs of simulated cohorts:

**Replicate** Two cohorts were simulated with the same parameter values, within the same paradigm

**Intra-paradigm** The two cohorts were simulated within the same paradigm, but with different parameter values (drawn from the same prior distributions)

**Inter-paradigm** The two cohorts were simulated in different paradigms, with different parameter values (randomly selected, and drawn from the same prior distributions if the parameter is shared across the two paradigms)

Note that the structure implies a fourth similarity level, with the same parameter values but differing paradigms. Due to the structure of this identifiability analysis, with a set number of parameter value sets randomly drawn within each paradigm for simulation, no pairs of simulations fall into this similarity level.

A shortcoming of this analysis is the lack of a proper treatment of subsampling: an area of future research must be to incorporate the effect of multiple random subsamples from each simulation, each of the number of cells sequenced from that patient, to investigate the effect that taking a single sample and only sequencing a fraction of the basal cells contained therein had on the identifiability of paradigms from the output. As presented in this work, subsampling is incorporated into distances using the phylogeny by taking a single subsample.

##### Replicate simulations show remarkable consistency

First, to assess the distance functions for their ability to detect change in parameter values from stochastic changes in simulations, we generated 100 different parameter sets in each hypothetical paradigm, and ran 2 replicate simulations with each parameter set.

These data showed remarkably small distances between replicate simulations relative to intra- and inter-paradigm distances across all distance functions (Figure S6A). This is reassuring: distance functions identified aspects of the data unlikely to be different at random. However, while in all cases aside from the negative control functions there is a significant difference both between replicate/non-replicate and between intra-/inter-paradigm distances (Mann-Whitney's U test with Bonferroni correction), there is no large visible distinction between intra- and inter-paradigm distances (indicating a significant but, on average, small effect). This does not necessarily indicate a lack of meaningful distinction between the models according to these metrics: instead, it could be that there is clear distinction between some or all paradigms, but this is masked by high variance along dimensions on which the paradigms are identical.

##### Distance function aggregation

A natural way to simplify a system with several different distance functions is to attempt to unify them into a single distance function, which conveys information from all distance functions on the proximity or otherwise of each pair of simulations. Simply summing the distance function values is inadequate due to differing scales, and normalising the distance functions (as before, say by dividing by mean replicate distance values or overall mean values) retains the issue of different functions having different effective weightings, due to their differing coefficients of variation.

To combine these distance functions in a way that equally weights variation in each distance function, we instead divided each distance value by the range between the $0.5^{th}$ and $99.5^{th}$ percentile of their distances, equalising the scales of each distance function in a manner robust to large outliers. This normalisation substantially reduced the discrepancy in standard deviation between the different distance functions, further than simple division by the mean value (Figures S6B–S6C).

We used this aggregated distance metric to create a simple k-nearest-neighbour (k-NN) classifier. This had limited accuracy in the problem of ascertaining the paradigm most likely to have generated an unseen simulation (Figure S7A).

**Redundant distance function elimination**

The distance functions as described above were highly correlated, unsurprisingly given that many are variations on similar themes. It is reasonable to question whether these are all necessary, or whether some subset of the distance functions may encode all or most of the information they contain - in particular, in the context of an approach based on aggregating the distance functions, any sets of equivalent distance functions would receive additional weight compared to other distance functions without providing additional information and thus reduce the effectively available information. To assess which distance functions provide the least additional information, we used the variance inflation factor (VIF) of each distance function's values across all pairs of simulated cohorts. This provides a measure of the degree to which each distance function's values over all pairs of simulated cohorts can be explained by a linear combination of other distance functions' values.

To find a subset of functions whose values provide meaningfully different information, we performed a greedy search to minimise the VIF of each distance function. At each iteration, VIFs were re-calculated for the new, smaller set of distance functions and the distance function with the highest VIF was removed. Iterations continued until a VIF threshold was reached by all distance functions: 5 is a standard VIF threshold in the different context of regression, so we used both this threshold and the looser threshold of 20. These correspond to 80% or 95% of a distance function's variance, respectively, being explainable by variance in other distance functions.

One distance function included (Wasserstein with smoking; see Table S4) is by definition a linear combination of other distance functions. It was therefore omitted from this analysis as it would lead to uninformatively infinite VIF values.

This analysis (shown in Figure S7C) revealed a core set of independent distance functions, and showed (via tracking of which distance functions reduce most in VIF after each removal) that while distance functions using the same modality are the most correlated, phylogenetic and mutational burden distributional distances are also closely related. For example, the removal of the 1-Wasserstein distance between mutational burden distributions had the strongest impact on the 1-Wasserstein distance between distributions of branch lengths in the phylogeny, despite their apparently differing measures. This would seem to indicate that, within this simulation model, mutational burden distribution is highly linked to the shape of the phylogenetic tree. This suggests that a degree of the information provided by the branch length Wasserstein comparison is in fact comparing mutational burden, due to the scaling of branch lengths to the number of mutations. An alternative scaling by the fraction of the mean number of mutations in the sample would remove this, and therefore perhaps provide a cleaner representation of information from the phylogeny. Table S7 shows the remaining distance functions after filtering with VIF thresholds 5 and 20.

This restricted set of distance functions provides a similar level of accuracy when summed as the unrestricted set (Figure S7B), not bringing the aimed-for improvement in identifiability. This analysis has significant limitations, relying as it does on only linear combinations and ignoring that some distance functions be significantly informative only for simulations in certain parameter regimes.

To improve this assessment of identifiability, we used a different approach to integrating the different metrics: embedding each in a Euclidean space and concatenating the spaces. This allowed for an orthogonal approach to identifying which distance functions are most useful.

##### Multi-dimensional scaling allows for training classifiers on simulation metrics

Classification is a standard task achieved in many machine learning applications, with a large body of practical research providing effective out-of-the-box packages capable of fitting, testing and benchmarking models. In this context, classifiers can be used to find a lower bound on the amount of information about the paradigm encoded in these metrics, applied to these simulations.

A powerful approach in answering a question such as “how much information about the simulation paradigm can be extracted from a simulation output via these metrics?” is to attempt, in some consistent and unbiased way, to learn the relationship between metrics and paradigms and check what accuracy is achieved on unseen samples. This approach, of using machine learning techniques to provide a lower bound on the identifiability of some inverse problem, has certain natural pitfalls (around over-fitting and accidental barcoding) to which true out-of-sample testing is a generally robust antidote.

This question almost falls neatly into the format of a classification problem, but for the fact that we were faced with a matrix (or indeed, several matrices, one for each distance function) of distances between our points, rather than a number of features (typically arranged in a vector) assigned to each simulation, which a classifier could train and test itself on. Unlike in the setting of convolutional neural networks for image analysis[72], where a matrix of values can meaningfully be viewed as a vector with some additional structure that can be encoded, in this case the matrix represents dissimilarities between objects.

In order to convert our distance matrix into features for each simulation, we used multi-dimensional scaling (MDS)[73], an established method to find vectors in N-dimensional space (for a given positive integer N), optimised to have the Euclidean distances between them as close as possible to the given values. We used this method, as implemented in the *sklearn.manifold* package in Python[44] using the SMACOF (Scaling by MAjorizing a COmplicated Function) iterative stress reduction algorithm of Kruskal[74,75], to create a vector of “features” of a specified length (values 2, 3, 5, 10 and 20 all tested) for each simulation, for each distance function. We then concatenated the vectors for each distance function-simulation pairing into a larger vector for each simulation, which could then be fed into a classifier with the generating paradigm as the label, to assess how well an un-tuned classifier could perform at learning the inverse relationship (mapping from simulation feature vector to paradigm) in cross-validation.

We leveraged the observation within the initial dataset of simulated cohorts that variation between replicates was small compared to that between differing simulations (Figure S6A) to increase the coverage of the prior parameter distributions by creating a larger dataset of 300 simulations in each paradigm, with only one replicate of each. Calculating all pairwise distances in this new set of simulated cohorts provided a larger set of distinct simulations, for which we generated features with MDS.

2-dimensional MDS embeddings (Figure S8A) reveal apparent differences in the geometries generated by these distance metrics: some appear mostly restricted to a single-dimensional subspace with a small number of outliers, while the z-value transformed 1-Wasserstein distance separates different groups of paradigms into distinct areas.

The MDS embedding process is in general unable to find points in a Euclidean space perfectly replicating distances observed in a higher-dimensional and/or non-Euclidean one. The process is one of optimisation to reduce the “stress”, a measure of the inaccuracy of the distance matrix. The final, minimised value of this stress can be used as a measure of the degree to which each distance function’s values were incompatible with projection into a lower dimensional space[73]. Those with particularly high optimal stress values are likely to have lost a significant amount of information in the embedding, and are thus expected to be less useful in the subsequent classification process. Figure S8B shows three notable trends in the optimal stress values:

- Higher dimensionality of the embedding space leads to lower stress. This is as expected: with more dimensions to vary, the MDS algorithm can improve its embedding.
- In cases where a distance function is repeated twice, once using the sum of squared values for each patient and once the sum of absolute values for each patient, the squared value iteration incurs higher stress.
- Distances associated with the means of the components of Gaussian mixture models have significantly higher stress than other distance functions. This seems to reflect a fundamentally non-Euclidean nature of these metrics, possibly due to discontinuities around the point where two peaks move past each other. These functions also seem to have higher stress values for higher dimension spaces, indicating problems with the optimisation algorithm.

These stress values provide a readout of the degree to which MDS failed to fully incorporate the dissimilarity information they encode. It may be that the information they successfully encode is all the useful information, and they failed to capture some useless noise that is irrelevant for the identifiability question, or it may be entirely the opposite (with mainly useful information thrown away in the MDS embedding and only useless noise retained): in a later section, we used the classification approach described in the remainder of this chapter to investigate which distance functions’ MDS features are most useful in this regard.

##### Classifiers trained on MDS embeddings of simulation outputs

Given a dataset of features and labels, the degree of information in the features pertaining to the labels can be ascertained via cross-validation, by splitting the data into 5 “folds”. Various algorithms have been designed and published to learn a relationship from the combination of 4 folds, which can then test their ability to predict the labels of the 5th, a process that can be repeated with each fold held out. As well as simple accuracy (the fraction of simulations assigned the correct unseen paradigm labels), models may provide information on the importance of each feature and the frequencies of each particular misclassification. Importance values can be averaged over the features contributed by each distance function to infer an importance score.

We used two standard classification methods in this study, to ensure that any accuracy is due to identifiability of the problem rather than a coincidental aptitude of a specific algorithm. The models were chosen due to their differences in approach, relative simplicity and ubiquity within modern machine learning approaches, and were applied "out of the box" as supplied by the *scikit.learn* package, to avoid over-fitting. The classification methodologies are briefly summarised below.

- Logistic regression, when used as a classifier, splits the problem into a separate binary prediction problem for each paradigm label. For each of these binary problems, a generalised linear model with logistic link function is fit, learning a linear relationship from the features to a probability that a given simulation will be from that paradigm. These probabilities can then be compared, and the highest-probability paradigm is predicted. This is equivalent to finding linear separating hyper-planes in the feature space between the different classes so as to optimise a cost function penalising ill-separated training datapoints. Importance of each feature can be inferred by normalising features before training the classifier and taking the absolute value of the coefficient assigned to each feature.
- Random Forest-based classification is a more modern method, which generates a large number of tree-based classifiers in a stochastic manner, each with high variance and access to only a subset of the data. These trees are then aggregated into an ensemble, which empirically performs well in many classification problems[76]. Importance of each feature is taken to be the Gini importance value as a first, computationally easy estimate: the manual approach shown in Figure 3C and described below is used for validation.

##### Support Vector Machine classifier omitted for low accuracy

The Support Vector Machine (SVM) architecture[77] was additionally used. This utilises the "kernel trick", whereby linear separation of classes is carried out in an implicit high-dimensional inner product space embedding of the data, only accessed via its images in the lower-dimensional feature space. The radial basis function was used as the kernel, a commonly used kernel in general biological applications[78] and the default option in the *scikit.learn* Python package[44]. Default values were taken for the kernel's hyper-parameters in order to assess how much information was readily available rather than find the best-possible fit.

This methodology showed significantly lower accuracy classifying unseen simulations with known ground truth (Figure S9A); as such it was dropped for downstream analysis.

##### Distance metric importance assignment

Aggregated feature importances over the features generated from each distance function allow an understanding to be gained of the degree of weight assigned to each distance function. Here, the negative control random distance provides a baseline level of importance generated by random noise. These importance scores can be considered either by raw value (normalised to have sum 1) or by their rank (Figure S10A), with the two measurements concurring on the ordering: z-value transformed 1-Wasserstein distance of mutational burden distributions is given a high score, along with other transformed mutational burden distribution metrics, the tree balance metric $J^1$ and measures of the size of the larger weight in Gaussian mixture models fitted to the mutational burden distribution (a measure of its degree of bimodality).

Other metrics are given lower scores: indeed, Figure S10A shows that they are even ranked on average below the random negative control.

As a result of known shortcomings of internal importance assignments, to verify that these importance scores accurately reflected the degree to which these distance functions were required for inference, we reran the classification with different subsets of the distance functions. For computational and combinatoric reasons, we considered only subsets of at most three distance functions, in order to assess each metric's ability to contribute to accuracy.

Comparing the cross-validation accuracy in classifiers with and without each individual distance function showed (Figure 3C) a broad concurrence with the classifier importance scores, with normalised 1-Wasserstein distances of mutational burden distributions, comparisons of phylogenetic trees and comparison of the bimodality in a Gaussian mixture model fit to the mutational burden distribution all improving cross-validation accuracy to a large extent, while other distance functions (those with a median rank below that of the random control in the importance score analysis) showed negative relative informativeness. This includes, reassuringly, the random control distance function.

This analysis, by restricting to small subsets of distributions, only considered the ability to contribute to a largely uninformed classifier. To consider the degree of added information over all other metrics, I repeated the analysis with each of these subsets of size at most three instead removed from the whole set. As there are eighteen metrics considered in total, this is equivalent to considering all subsets of size at least fifteen. While the previous analysis considered which distance functions were relatively more helpful than others, this approach instead sheds light on the distinct question of which functions provide additional information given knowledge of the values of every other function, analogous to forwards and backwards variable selection.

Note that negative values here reflect a relative rather than absolute lack of utility: due to the limited size of subsets considered, inclusion of different distance functions are not independent and to include one necessarily reduces the likelihood of inclusion of another distance function. Due to the unbiased inclusion of all subsets at these sizes, the ranking of the distance functions within each group is valid.

##### Restricting training simulations

A parameter of this identifiability analysis which was chosen entirely for computational reasons was the number of simulations run in each paradigm. To test that this is sufficient, and that those paradigms labelled as harder to distinguish were not simply those for which the analysis was underpowered, we reran the classifiers with smaller sets of simulated cohorts to train on. Out of the total 300 simulations in each paradigm, subsets of sizes 5, 10, 20, 50, 100, 150 and 200 were drawn (3 subsets for each size), and classifiers retrained on only the MDS features from those subsets. As the MDS embedding was a significant bottleneck, this was not rerun with only the subsetted distances; it is possible that some information may leak into the MDS features of a subset of simulations from those with which it was embedded, even without including those additional simulations in the training or testing sets. Nonetheless, this analysis allowed inference of the relative effect of increasing and decreasing sizes of training datasets.

Classifier accuracy was indeed dependent on the size of the simulation subset (Figure S10C), broadly consistently across all classification methods. There was also a notable increase in the random variance of accuracy at lower values (Figure S10D), as the classifiers become closer to random guesses. As well as being less accurate than the other classification methods generally, the support vector machine is also more sensitive to reduction in simulation count, with performance approximately random at the lowest value. Saturation in cross-validation accuracy seemed to occur well below 300 simulations per paradigm, justifying this size of dataset of simulated cohorts in the trade-off between accuracy and computational cost.

##### Restricting patient cohort

Another question this analysis can answer is one of the effect of cohort size: does this level of identifiability of outputs from these metrics applied to simulations remain if only a subset of the patient cohort is included in these distances? We reran all classifiers on these reduced distance functions for each of four patient groupings:

**Total** The whole cohort, for ease of comparison

**Yoshida et al. Patients** Only those patients from the Yoshida et al. cohort.

**Status Representative Patients** Only three patients included: the single smoker, ex-smoker and never-smoker with the most cells sequenced out of their smoking status.

**Huang et al. Patients** Only those patients whose lung stem cells were sequenced by Huang et al[4]. Note that since the identifiability analysis does not assess the effect of subsampling in any other than phylogenetic distances, the fact that this cohort had fewer cells sequenced per patient has a reduced impact on identifiability.

Figure S10E shows cross-validation accuracy decreases down this list of subsets, with the Yoshida et al. cohort providing accuracy levels nearly comparable to the total cohort, and three patients from this study marginally winning out over the Huang et al. cohort. However, these effects are small: as shown in the plot, the effect of choosing a different number of dimensions for each distance function's MDS embedding has a greater effect.

This is somewhat surprising, and may reflect a limitation of this simulation study rather than a saturation in cohort size: as we intended to use this small cohort of patients to discern the likeliest mechanism at work in the human lung in general (rather than in each patient's individual lungs), distinct from the admirable and important trend of personalised medicine, the simulation model laid out above is only marginally dependent on the patients' personal histories. It is therefore perhaps not surprising that only a small number of patients is needed to pin down the dynamics at work. Incorporation of patient heterogeneity, either through differing environmental exposures, germline and sex differences or infrequent random events occurring over the course of life, is an exciting area for future progression in this area. It also (of course) brings significant additional challenges, and therefore falls outside the scope of this work. An initial change would be to consider replicate simulations at the level of each individual patient rather than the whole cohort, allowing rare random events to be better replicated.

#### Identifying and removing distance functions that measure true data outside the distribution of simulations

Any classification approach is unlikely to extend successfully to unexplored regions of its feature space. For the problem of extending the classifiers trained on simulations to the observed scWGS dataset, the feature space is the product of the implicit space created by each of a set of distance functions. To ensure that the true data is not in an unexplored region requires removing those distance functions with respect to which the true data is an outlier, without any close neighbours. At a philosophical level, this reflects the removal of those distance functions which, regardless of their ability to gather useful information about the simulations from their outputs, measure modalities of the data along which simulations are unable to recapitulate true data. Any simulation study is inevitably a compromise between parsimony and accurate reproduction of dynamics; this filtering removes those comparisons that overly focus on the simplifications over the reproduced underlying dynamics.

To determine which distance functions warranted exclusion, we calculated the distance from the true data to each simulated cohort by each distance function, and compared the distribution of each distance function's true-vs-simulation distances to that of its between-simulation distances (Figure 4A). For a given percentile threshold n, we then removed those distance functions for which the $n^{th}$ lowest percentile of the true-vs-simulation distances was higher than the $(100 - n)^{th}$ percentile of the between-simulation distances. According to distance functions removed by this filter, the true data is further away from almost all simulations than the total span of the simulations. Figure S8A shows 2-dimensional MDS projections of all distance functions, in increasing order of threshold required to exclude them. This removal had minimal effect on the importance ordering assigned to classifiers (Figures S11A–S11C).

#### All-paradigm classification

While MDS embedding-based classifiers showed relatively high accuracy at the level of small clusters among the restricted group of 9 "protection-selection" hypothesis combinations (see above), the harder problem of classifying a simulation among all 24 combinations has a reduced accuracy (Figure S9A), revealing a high identifiability of the quiescent subpopulation (Figure S9B), with limited accuracy in classifying other hypotheses.

Applying these broader classifiers to the observed data's embedding vectors shows a consistent selection of the quiescent subpopulation after filtering distance functions for observed data extremity (Figure S12D). While there is not direct indication in favour of the immune hypothesis in these lower-accuracy classifiers, they do not rule it out as they do the smoking driver hypothesis. This indicates that the classifier framework described in this work has limited accuracy in the less constrained problem of classifying all combinations of hypotheses, but the result reached by the more accurate constrained classifiers is not inconsistent with that of the all-paradigm classifiers when applied to the observed scWGS dataset.

#### Immune death rate titration

In order to select the prior mean value for the immune death rate, simulations were run with other parameters taken at default values, with the immune death rate varied. This then provides a means of assessing the empirical levels of immune cell turnover produced by different values of the parameter. From this analysis, the value 0.00025 was selected to provide a turnover of approximately $\frac{1}{4} \times 1.0\ \mathrm{cell}^{-1}\ \mathrm{year}^{-1}$ at age 40, the approximate age of patients assessed by Teixeira et al[23]. The immune death rate at this value is shown in Figure 2G.

#### Quiescent gland fixation probability

In order to model a quiescent gland as a single evolutionary unit with a single effective genome, a filter must be applied to new mutations checking whether the mutation reaches fixation in the gland or is lost after its inception. Basic assumptions are made about the small population of stem cells within a submucosal gland in order to make this problem tractable:

- The gland is small, with no meaningful spatial separation between any pair of stem cells.
- The number of stem cells maintaining the gland is constant over time, with differentiations and deaths quickly counteracted by symmetric divisions, as in the main epithelium.

Given these assumptions, a standard result of evolutionary theory (of Ewens[79], used here as stated by Parsons and Quince[80]) can be applied: given a population of N cells, of which M have type X with division and death rates $\lambda_X, \mu_X$ and the remainder are of type Y with corresponding rates, then in a non-spatial Moran model the probability that cell type X reaches fixation is

$$\mathrm{P(fixation)} = \frac{\nu^N - \nu^{N-M}}{\nu^N - 1}, \text{ where } \nu = \frac{\mu_Y \lambda_X}{\mu_X \lambda_Y}.$$

This formula is applied with $M = 1$ corresponding to a new mutation, using the relative division rates according to the fitness change f associated with the mutation:

$$\begin{aligned}
\lambda_X &= r + S_r(f) \\
\lambda_Y &= r \\
\mu_X &= r - S_r(f) \\
\mu_Y &= r,
\end{aligned}$$

where r is the symmetric division probability and $S_r$ is the sigmoid projection function defined in Methods. Rearranging, this gives $\nu = e^f$ and

$$\mathrm{P(fixation)} = \frac{e^{f(N-1)}\left(e^f - 1\right)}{e^{fN} - 1},$$

which for large Nf can be closely approximated by $1 - e^{-f}$. For each new ambient (i.e. not while dividing to create a main population basal cell) mutation arising in a quiescent gland, the mutation is discarded with probability $1 - P(\text{fixation})$.

#### Total patient-years simulated

This work uses the current understanding of evolutionary dynamics in human upper airway stem cell populations to draw further inference from genomic recovery data. This requires running experiments *in silico* of the effect of decades of tobacco exposure under different tissue structures. To demonstrate that this could not feasibly be carried out in any experimental models, we calculated the total number of patient-years simulated using our framework for this work. Additional simulations used for optimisation, likely to be a larger number than those used for final inference, are excluded.

First, each simulation of the example smoking histories (as shown in Figures 2C–2D) simulates 240 patient-years. In each of the spatial and non-spatial regimes, for each of 24 hypothesis combinations, 10 replicate simulations were run, as well as an additional 10,000-cell simulation and 100 replicates in restricted hypothesis combinations in the spatial setting for Figures 2E–2H. An additional grid-search performed for calibration of parameters for the distribution of fitness effects adds $4 \times 14$ simulations. These simulations therefore in total took

$$240 \times (24 \times 11 \times 2 + 9 * 100 + 4 \times 14) = 356,160 \text{ patient-years.}$$

Simulations of the full cohort take 2818.416 patient-years (Figure 1A). The initial simulated cohort dataset, with 2 replicates of each of 100 simulations in each combination of hypotheses (shown in Figure S6A), took 7200 simulations. An additional 7200 were used for the subsequent repeat with 300 1-replicate simulations (shown in Figures 3, 4). Finally, to assess the parameter values best fitting the true data, we ran 10,000 full cohort simulations. Thus, full cohort simulations took in total

$$2818.416 \times (10,000 + 7200 + 7200) = 68,769,350.4$$

Thus, in total 69.1 million patient-years of simulations were run for the final results shown in this work. In vivo experiments with equivalent power run in parallel would require lifelong tracking of the entire population of Fiji, and run in series would need to have been run since the late Cretaceous, highlighting the necessity for computational approaches to address this question.

#### Supplemental Figures

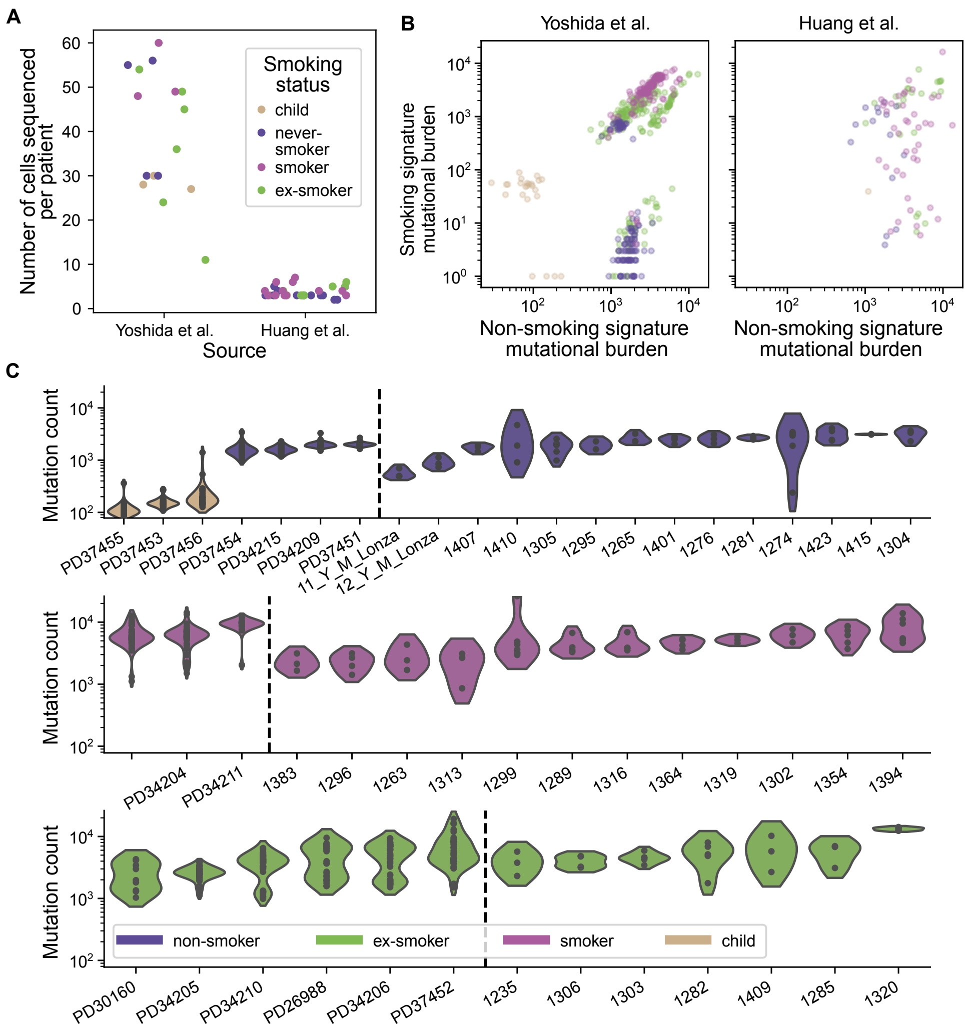


##### Figure S1. Further characteristics of combined cohort of scWGS data

(A) Number of cells sequenced via *in vitro* culture and bulk sequencing, from each patient stratified by cohort and coloured by smoking status as in Panel C.

(B) Number of mutations attributable to smoking and not attributable to smoking via signature deconvolution (assigned to SBS4, SBS92, DBS2 or ID3), within each cell sequenced in the combined cohort, coloured by smoking status as in Panel C and separated by study.

(C) Total number of somatic mutations in each sequenced basal cell, with kernel density estimates (generated by the *seaborn* package[16] using Scott's method for bandwidth calculation[17], with a cutoff of 1 bandwidth past extreme datapoints) to show the mutational burden distribution within each patient.

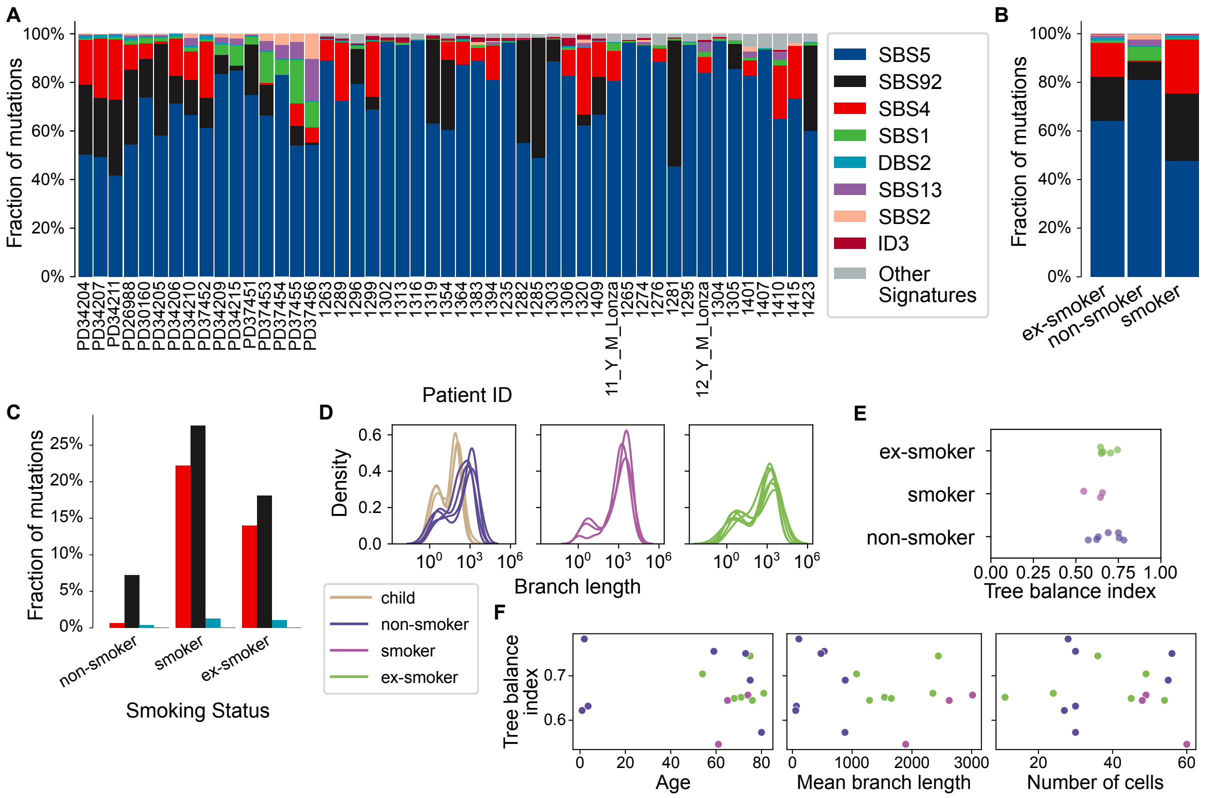

#### Figure S2. Signature deconvolution and phylogenetic summaries of the combined cohort of scWGS data

(A) Fractions of mutations attributable to each mutational signature within cells sequenced from each patient. Signatures with fewer than 0.5% total are combined as “Other signatures”.

(B) Fraction of mutations attributable to each mutational signature, aggregated by smoking status.

(C) Fraction of mutations attributable to each smoking-attributable mutational signature (SBS4, SBS92, DBS2 and ID3), aggregated by smoking status. Colours as in Panels A and B.

(D) Kernel density estimates (generated by the *seaborn* package[16] using Scott’s method for bandwidth calculation[17]) of the distribution of branch lengths within the phylogenetic trees inferred for each patient in the Yoshida et al. cohort, coloured by smoking status.

(E) Tree balance index for each patient in the Yoshida et al. cohort, by smoking status.

(F) Relation of tree balance index $J_1$ to patient age (left), mean phylogenetic branch length (center) and number of cells sequenced (right). All relations are statistically insignificant (Spearman’s rank-order correlation test, p-values 0.965, 0.770 and 0.892 respectively, to 3 significant figures).

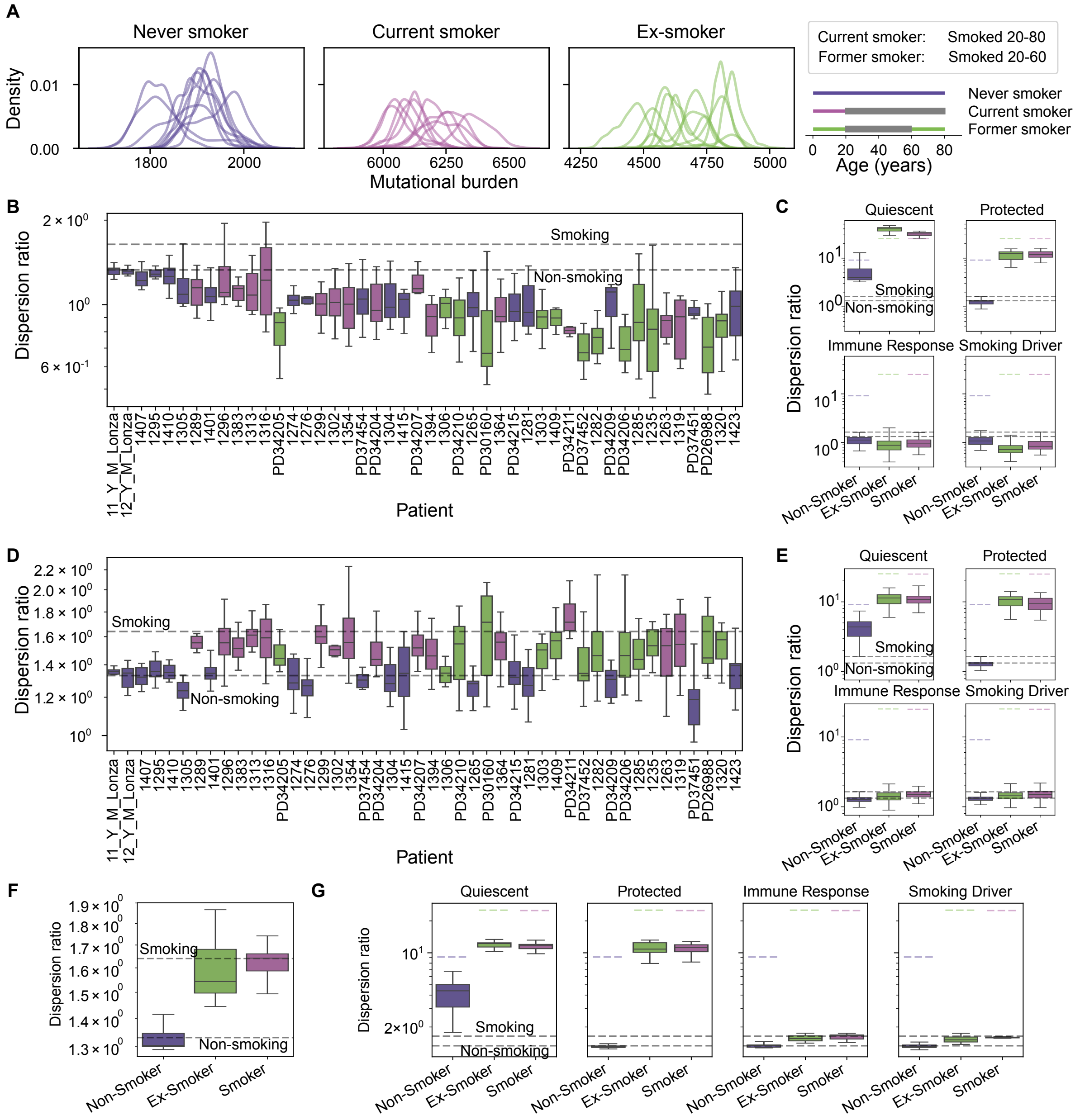


#### Figure S3. Non-spatial simulations fail to show sufficient variability in mutational burden; spatial simulations remedy this

(A) Kernel density estimates (generated by the *seaborn* package[16] using Scott's method for bandwidth calculation[17]) of distributions of mutational burden among simulated populations of airway basal cells for three example smoking statuses (as in Figures 2C–2D), simulated non-spatially over 10 replicates (separate lines) in the base paradigm with prior mean parameter values.

(B) Dispersion ratios (standard deviation as a fraction of the square root of the mean, as defined in Supplementary section "Theoretical dispersion") of mutational burden distribution in non-spatial simulations with no hypotheses activated of each patient in the cohort, coloured by smoking status. Dashed lines show expected value for the dispersion ratio under a model excluding cell-cell interaction, during smoking (above) or non-smoking (below). Boxplots: central line represents median, box indicates $25^{\text{th}}$ and $75^{\text{th}}$ percentiles, and whiskers indicate the minimum and maximum value after exclusion of outliers (shown as circles), determined by a cutoff of $1.5\times$ interquartile range outside the box.

(C) Dispersion ratios (as in Panel B) of mutational burden distribution by smoking status in non-spatial simulations of the combined cohort, with each individual hypothesis activated. Grey dashed lines and boxplots as in Panel B. Coloured dashed lines show the mean dispersion ratio of mutational burden of patients in the observed cohort, grouped by smoking status.

(D) As in panel B, with spatial simulations.

(E) As in panel C, with spatial simulations.

(F) Dispersion ratios (as in Panel B) by smoking status in 10,000-cell spatial simulations with no hypotheses activated of the combined cohort.

(G) As in E, with 10,000-cell spatial simulations.

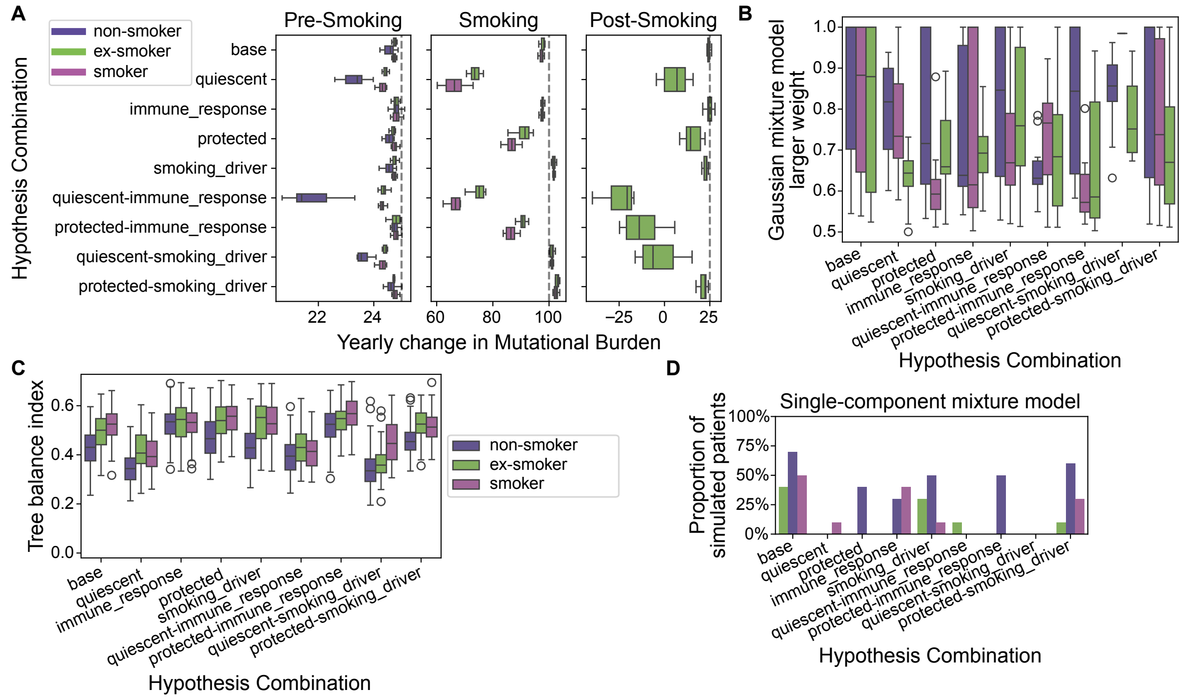


#### Figure S4. Summary statistics of simulations of example smoking histories, run with prior mean parameter values

(A) Mean yearly change in mutational burden in 10 replicate simulations of each smoking status example (as in Figures 2C–2D) with prior mean parameter values (see Supplementary section "Simulation parameters and prior distributions", Table S3), separated by the three stages of a smoking history, and the paradigm (Table S2) in which the simulation was run. Dashed lines indicate the rate of accumulation of individual cells. Boxplots: central line represents median, box indicates 25th and 75th percentiles, and whiskers indicate the minimum and maximum value after exclusion of outliers (shown as circles), determined by a cutoff of 1.5$\times$ interquartile range outside the box.

(B) Larger weight of a 1- or 2-component Gaussian mixture model fit to simulated (as in Panel A) mutational burden distributions, a measure of bimodality (0.5 denotes two peaks of equal weight).

(C) Tree balance index $J_1$ of phylogenies of simulated (as in Panel A) cell populations, coloured by smoking status. Boxplots as in Panel A.

(D) Proportion of simulated (as in Panel A) patients with a single-component Gaussian mixture model selected by BIC over a two-component model.

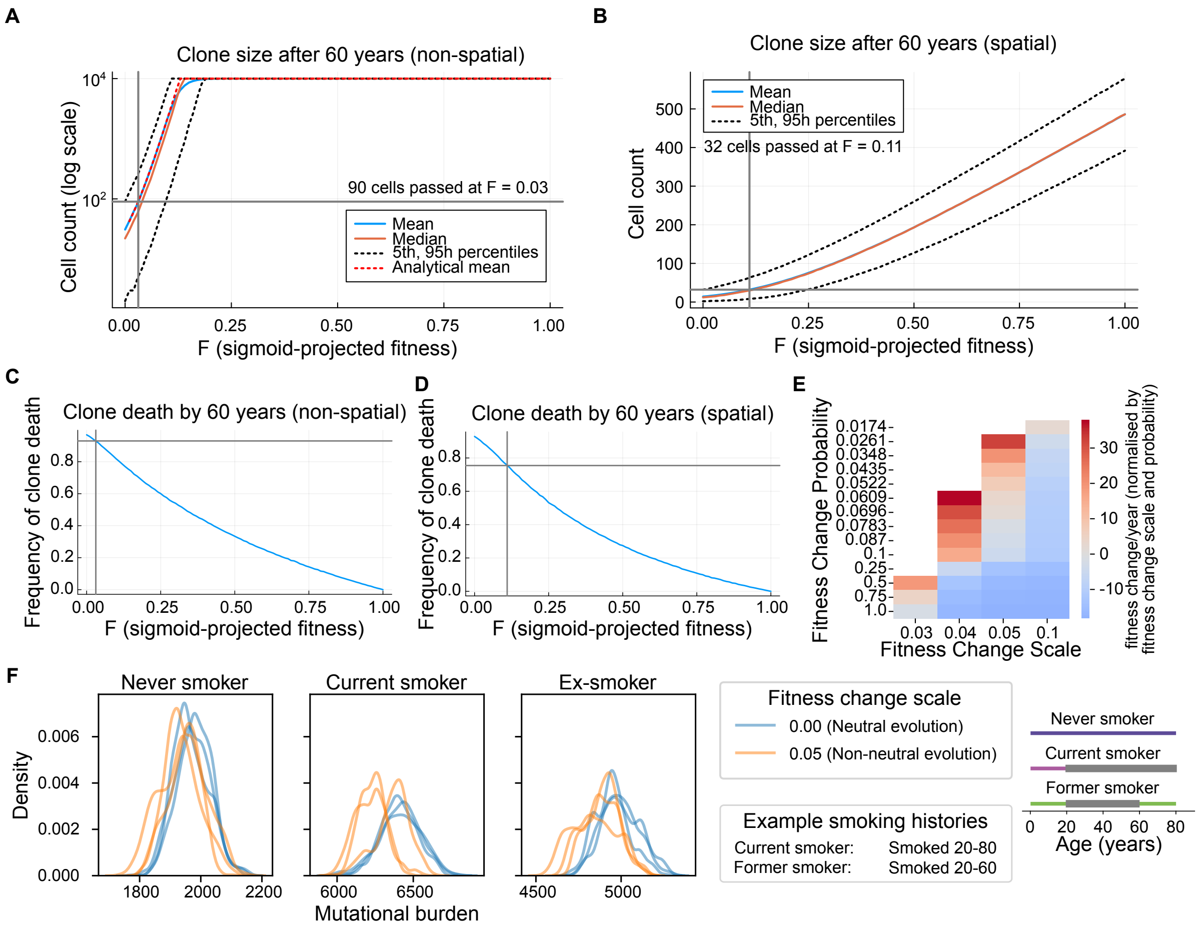


#### Figure S5. Inference of DFE for use in simulations via simplified simulation scheme

(A) Mean (blue line) and 5th and 95th percentile (dashed lines) of the empirical clone size after 60 years over $10^5$ Gillespie non-spatial (well-mixed) simulations of the continuous-time Markov Chain model for expansion of a single clone (see Supplementary section "Distribution of fitness effects") for each of a range of values of projected fitness F := $\sigma$(f). Simulations in which the clone reaches size 0 are excluded. A threshold for a mutation to be considered a significant driver is found by the level of F (0.03 in this case) at which the mean clone size after 60 years exceeds the $95^{th}$ percentile (90 cells in this case) in the neutral case F = 0. A cap of $10^4$ mutations was used for computational reasons; this does not impact calculation of the threshold.
(B) As in Panel A, with spatial Gillespie simulations. Here, the threshold is found at F = 0.11 to exceed the 32-cell threshold of significant difference from the neutral case. There is no analytical solution known to the authors in the spatial setting.
(C) Fraction of $10^5$ simulations (as in A) in which the clone reaches size 0 (an absorbing state) before 60 years in simplified non-spatial simulations of expansion of a single clone (see Supplementary section "Distribution of fitness effects"), by initial projected fitness value F. The threshold F = 0.03 found in Panel A is shown, at which 93.0% of simulated clones reached size 0 before 60 years.
(D) As in Panel C, with spatial simulations. The threshold F = 0.11 is shown, at which 75.4% of simulated clones reached size 0 before age 60.
(E) Mean yearly fitness change (normalised by fitness change scale and fitness change probability, noting sign is unchanged by this) in spatial simulations with different fitness change scale and fitness change probability values. Values incompatible with observed driver frequencies (see Supplementary section "Distribution of fitness effects") were not simulated and are left blank. Values for fitness change probability were chosen based on round fractions of unity and round multiples of 0.0174, the exonic fraction of the genome.
(F) Kernel density estimates (generated by the *seaborn* package[16] using Scott's method for bandwidth calculation[17]) of distributions of mutational burden in simulations of example smoking histories (as in Figures 2C–2D) with fitness included (at the selected values 0.1, 0.05 for fitness change probability and fitness change scale respectively) or excluded (by setting the fitness change scale scale to 0).

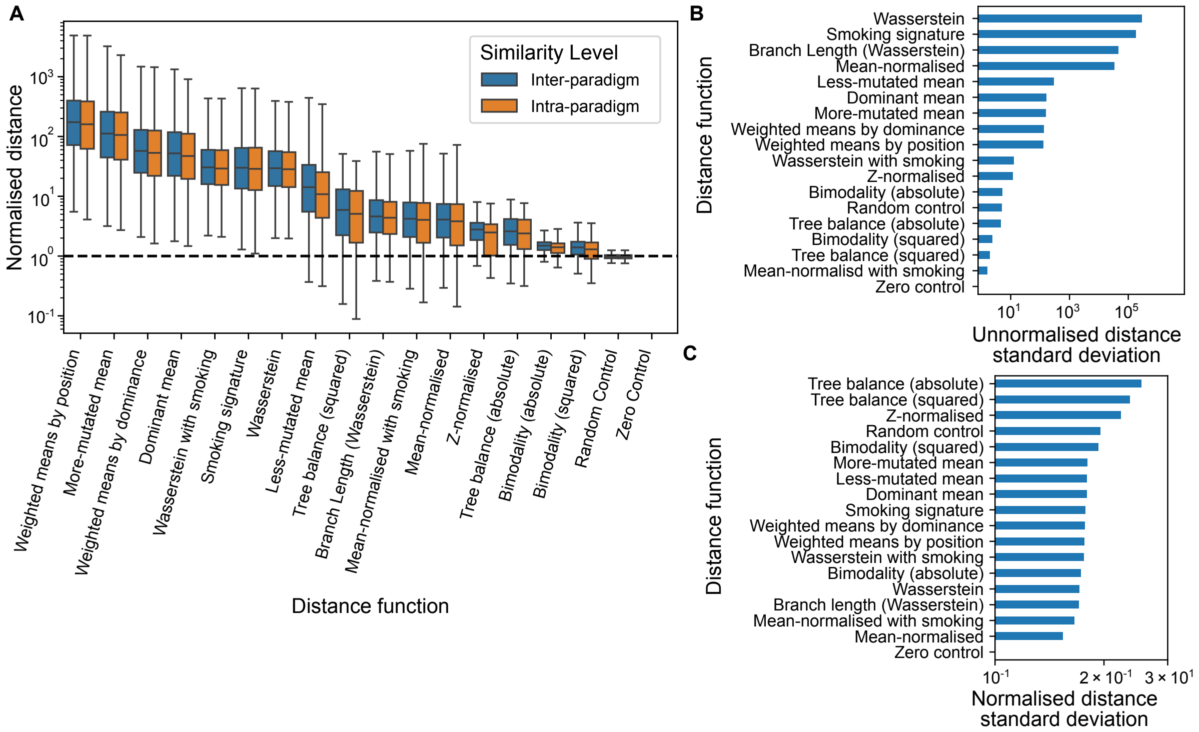


**Figure S6. Simulations are more different than replicates, and distances can be normalised to reduce variance**

(A) Distance between pairs of simulations run with different (inter-paradigm) or the same (intra-paradigm) set of hypotheses activated, for each distance metric used, in a dataset of 100 simulations per paradigm, each run for 2 replicates (see Methods). Distances are normalised by the mean distance between replicate simulations, by that metric. Boxplots: central line represents median, box indicates 25$^{th}$ and 75$^{th}$ percentiles, and whiskers indicate the minimum and maximum value after exclusion of outliers (shown as circles), determined by a cutoff of 1.5$\times$ interquartile range outside the box.

(B) Sample standard deviation of the set of all unnormalised pairwise distances between simulations in a dataset of 300 single-replicate simulations per paradigm (see Methods).

(C) As in Panel B, after normalising by the difference between the 0.5$^{th}$ and 99.5$^{th}$ percentiles of that distance function's values.

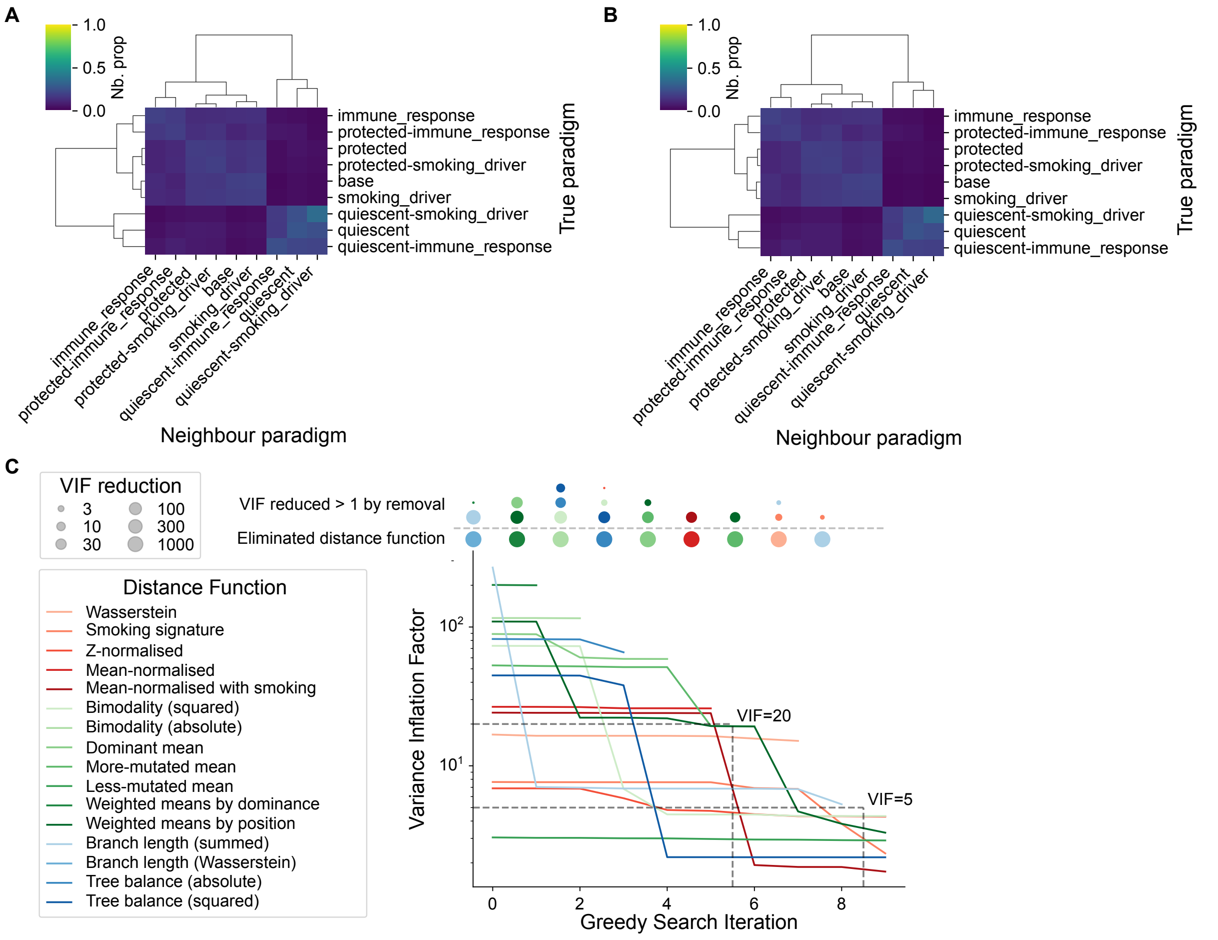


#### Figure S7. Filtering distance metrics to a minimal independent set does not improve aggregated distance classification

(A) Accuracy matrix plot (as in Figure 3B) for a k-nearest-neighbour classifier using the full aggregated distance, showing the proportion of nearest 5% of neighbours of each simulation run in each paradigm (row) which belongs to each other paradigm (column).

(B) As in Panel A, after filtering out distance functions via the greedy search VIF minimisation algorithm with threshold 5.

(C) Distance functions removed at each stage of the greedy search VIF minimisation algorithm to find an independent set of distance metrics. The distance function removed, and those distance functions whose VIF is significantly reduced by this, are shown for each step. Lines show linear interpolation between distance functions' VIF values at each iteration.

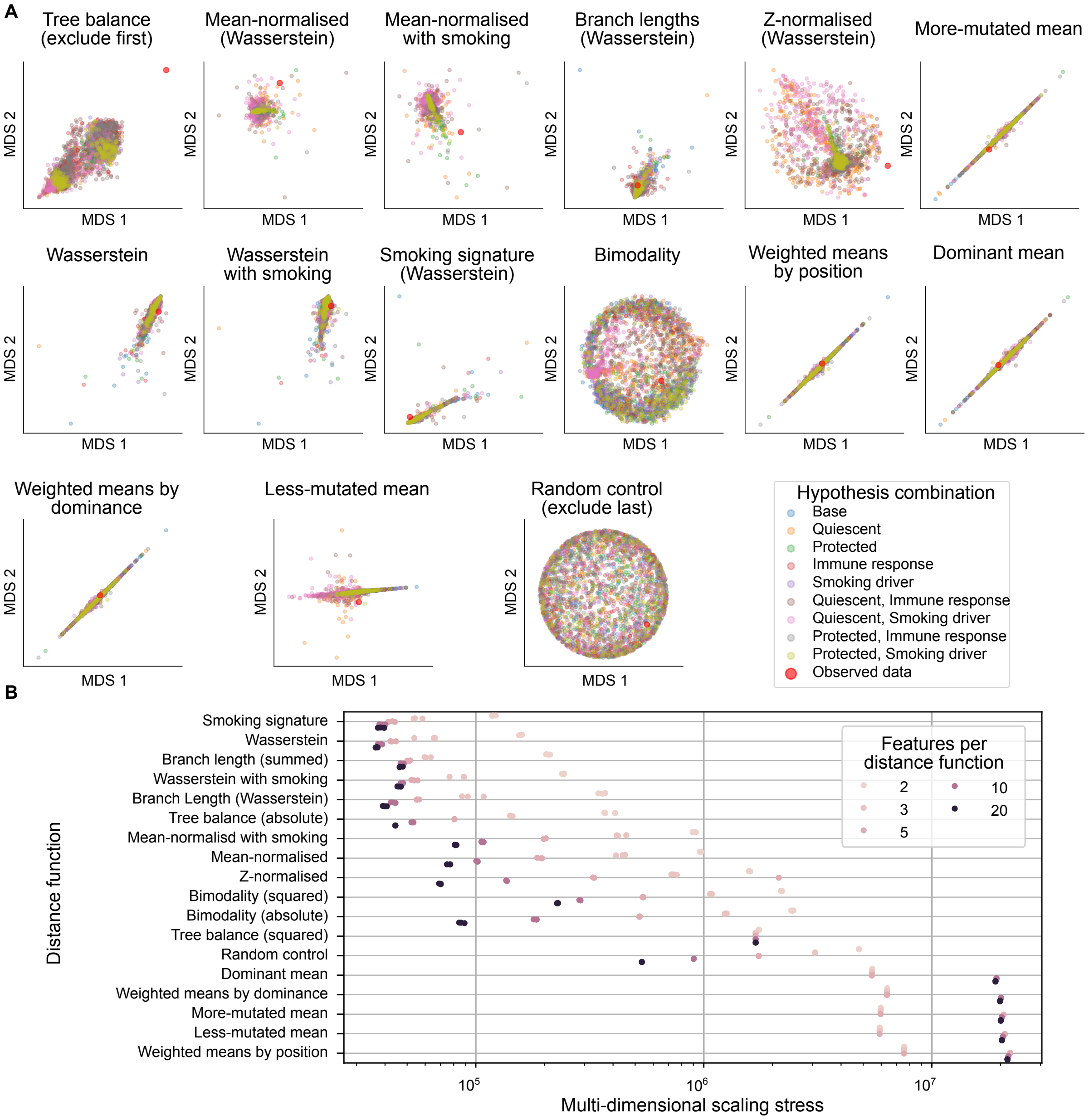


**Figure S8. Multi-dimensional scaling embedding of simulated cohorts with respect to different distance metrics, including MDS projections for distance functions first eliminated for true data extremity**

(A) Two-dimensional MDS embedding of pairwise distances between simulated cohorts (coloured by paradigm), for each distance metric described in Tables S4–S6. Each point represents a simulated cohort.

(B) Stress incurred[74] in MDS embedding of each distance function, given different dimensionalities of embedding.

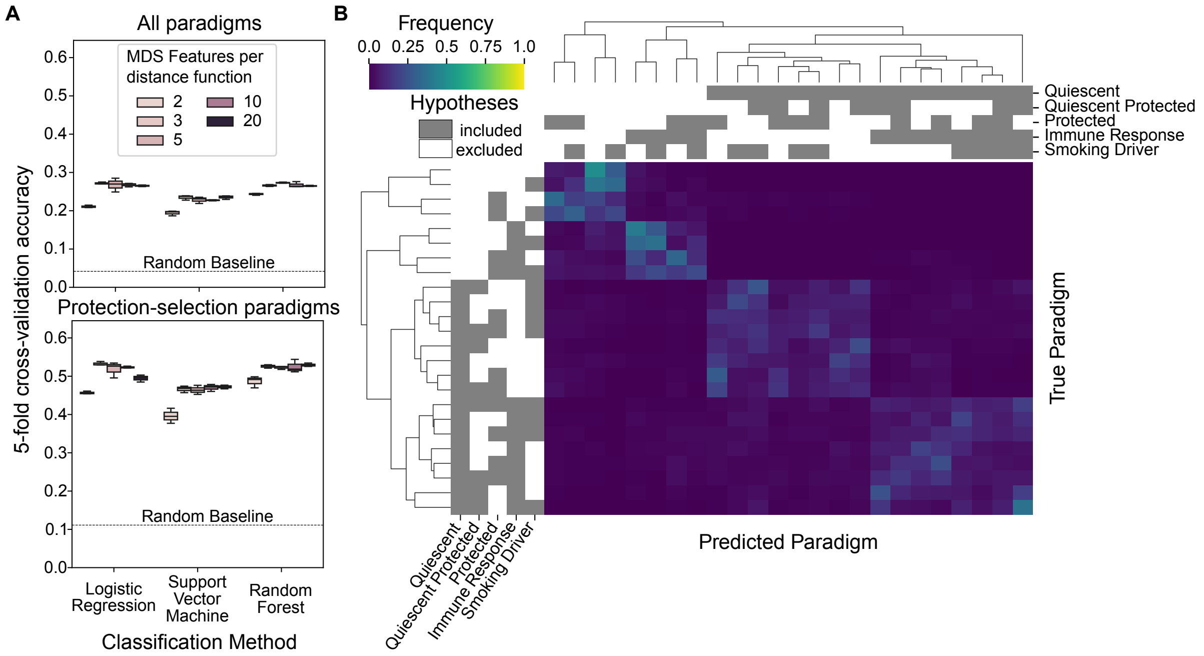


**Figure S9. Classifier identifiability analysis applied to all combinations of hypotheses**
(A) 5-fold cross-validation accuracy of classifiers trained on concatenated MDS embeddings of all distance metrics on all 24 paradigms (above) or the restricted set of 9 “protection-selection” paradigms (below). Boxplots: central line represents median, box indicates 25$^{th}$ and 75$^{th}$ percentiles, and whiskers indicate the minimum and maximum value.
(B) Mean classification frequency across all classifiers of each paradigm, in assessing outputs from unseen simulations with each other ’True’ paradigm, for the all-paradigm classification problem.

| Patient ID | Silverman, 1981 | Hall and York, 2001 | Fisher and Marron, 2001 | Hartigan and Hartigan, 1985 | Cheng and Hall,1998 | Ameijeiras-Alonso et al, 2019 |
|---|---|---|---|---|---|---|
| PD26988 | 0.033280 | 0.034080 | 0.033547 | 0.033653 | 0.032640 | 0.032320 |
| PD30160 | 0.299264 | 0.295008 | 0.296000 | 0.295424 | 0.297824 | 0.296928 |
| PD34204 | 0.299264 | 0.295008 | 0.296000 | 0.295424 | 0.297824 | 0.296928 |
| PD34205 | 0.865600 | 0.898978 | 0.834149 | 0.852086 | 0.843815 | 0.900689 |
| PD34206 | 0.000000 | 0.000000 | 0.000000 | 0.000160 | 0.000480 | 0.000480 |
| PD34207 | 0.979270 | 0.980770 | 0.978710 | 0.979230 | 0.979160 | 0.981520 |
| PD34209 | 0.865600 | 0.898978 | 0.834149 | 0.852086 | 0.843815 | 0.900689 |
| PD34210 | 0.001520 | 0.000960 | 0.001280 | 0.001120 | 0.000960 | 0.001120 |
| PD34211 | 0.865600 | 0.898978 | 0.834149 | 0.852086 | 0.843815 | 0.900689 |
| PD34215 | 0.667520 | 0.638507 | 0.667307 | 0.645173 | 0.610027 | 0.725333 |
| PD37451 | 0.865600 | 0.898978 | 0.834149 | 0.852086 | 0.843815 | 0.900689 |
| PD37452 | 0.865600 | 0.898978 | 0.834149 | 0.852086 | 0.843815 | 0.900689 |
| PD37453 | 0.979270 | 0.980770 | 0.930677 | 0.979230 | 0.961067 | 0.981520 |
| PD37454 | 0.882194 | 0.898978 | 0.834149 | 0.868194 | 0.866103 | 0.900689 |
| PD37455 | 0.865600 | 0.901589 | 0.834149 | 0.852086 | 0.843815 | 0.907029 |
| PD37456 | 0.865600 | 0.898978 | 0.834149 | 0.852086 | 0.843815 | 0.900689 |

**Table S1. P-values for tests for bimodality of mutational burden distributions of each patient** Benjamini-Hochberg- adjusted p-values calculated under six distinct statistical tests for bimodality[38–43], applied to the distribution of mutational burden within the sampled cells of each patient in the combined cohort[3,4]. The *multimode* R package[43] was used, with a bootstrap replicate value of $10^5$.

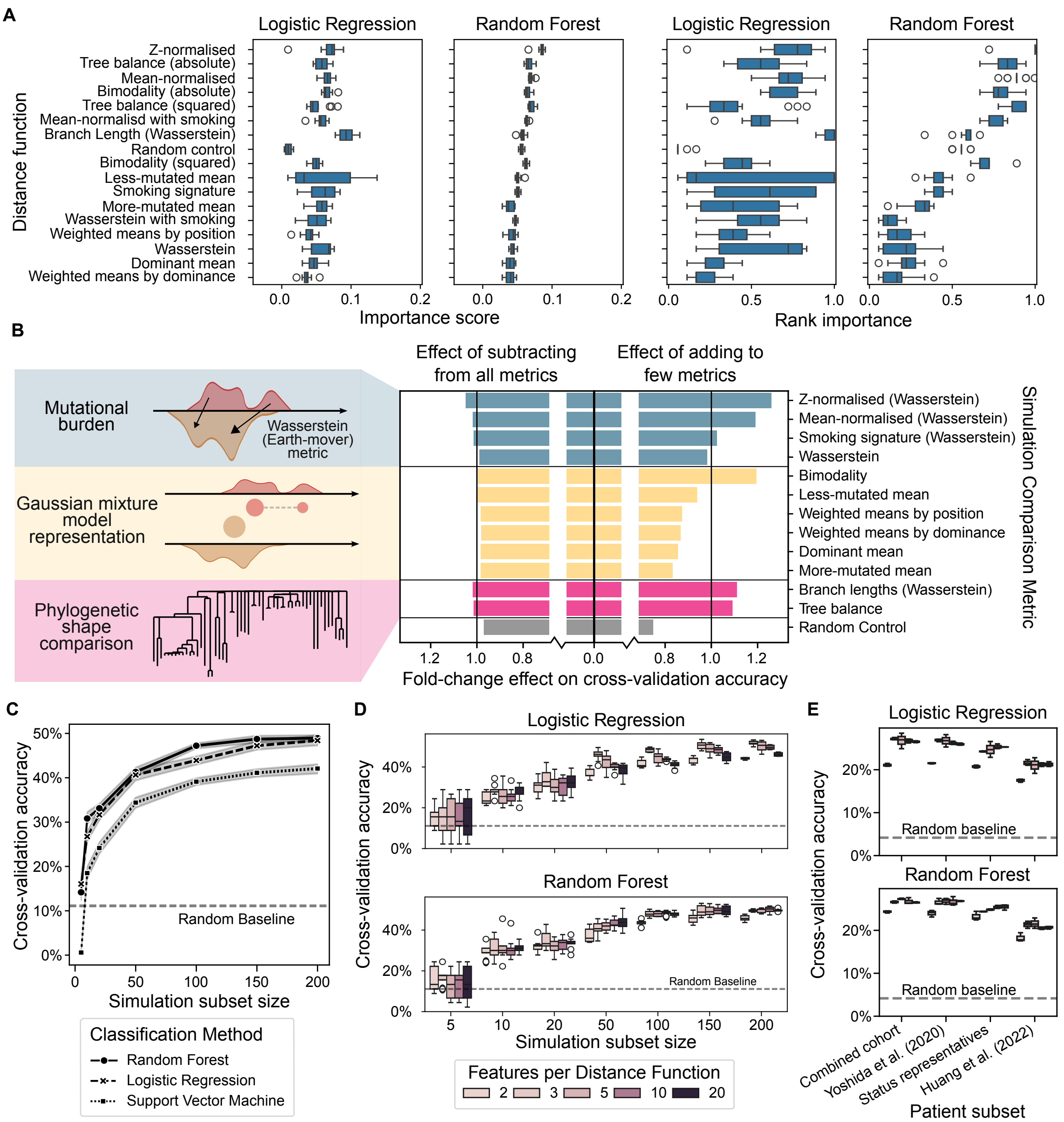


#### Figure S10. Restricting classifiers reduces their accuracy

(A) Classifier-intrinsic mean importance score over features derived from each distance metric's MDS embedding. Raw importance score (left) and rank importance (right, see Methods) shown for each classification methodology. Boxplots: central line represents median, box indicates 25th and 75th percentiles, and whiskers indicate the minimum and maximum value after exclusion of outliers (shown as circles), determined by a cutoff of 1.5× interquartile range outside the box.

(B) Impact on accuracy of each biologically-motivated metric used to train models on simulation outputs. Fold change effect on cross-validation accuracy is shown of inclusion vs exclusion of each distance function within all subsets of size at most 3 (right) and at least 14 (left).

(C) Cross-validation accuracy of classifiers trained on subsets of simulations of different size. Simulation subset size refers to the number of simulations within each paradigm. Points show mean values over 3 replicate subsamplings of simulation cohort at each size, shaded areas show a bootstrapped 95% CI and lines shown linear interpolation.

(D) Cross-validation accuracy by embedding feature dimensionality and training subset size. Boxplots as in Panel A; simulation subsets as in Panel C.

(E) Cross-validation accuracy when trained on different restrictions of the patient cohort (see Supplementary section "Restricting patient cohort").

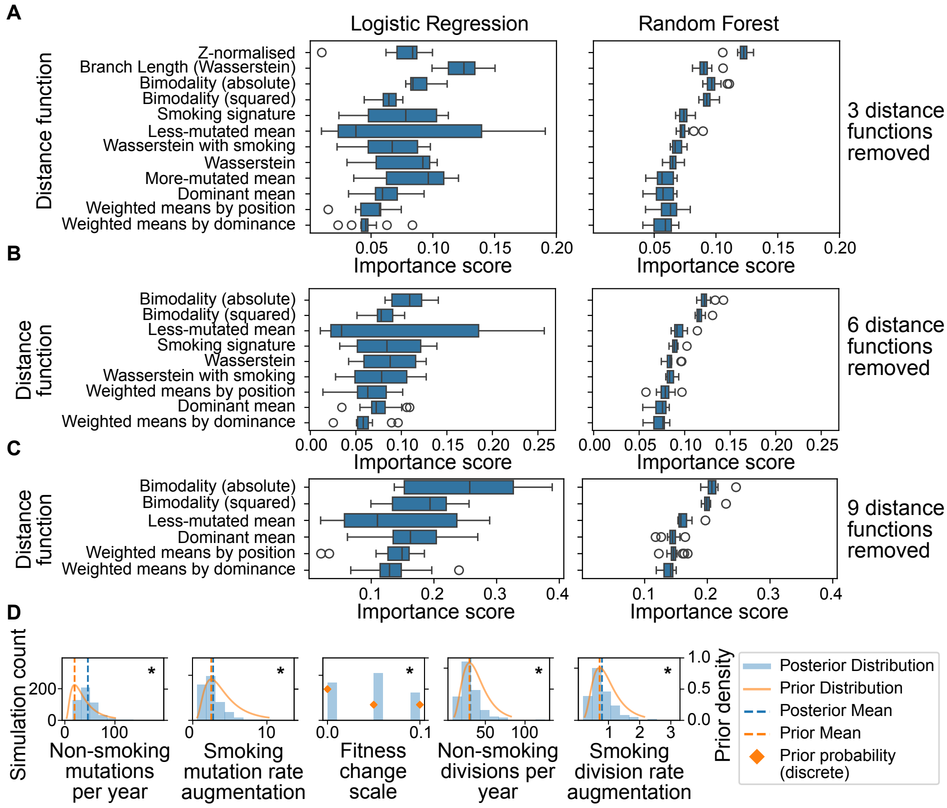


#### Figure S11. Importance of distance functions in classifiers after filtering for observed data extremity

(A) Mean importance score over features derived from each distance function, in classifiers after removal of three metrics for observed data extremity. Boxplots: central line represents median, box indicates 25$^{th}$ and 75$^{th}$ percentiles, and whiskers indicate the minimum and maximum value after exclusion of outliers (shown as circles), determined by a cutoff of 1.5$\times$ interquartile range outside the box.

(B) As (A) after removal of 6 metrics.

(C) As (A) after removal of 9 metrics.

(D) Distribution of parameters not shown in Figure 4F, in 5% of simulations most closely approximating (by summed normalised distances) the observed dataset, within the quiescent and immune response paradigm, at the most stringent observed data extremity threshold. 500 simulations shown out of 10,000 run with parameters drawn from prior distributions (shown in yellow, as probability density or mass functions; see Table S3). Prior and posterior means are shown. Asterisks denote parameters for which the posterior distribution is significantly different to the prior distribution (see Table S8 for p-values and a full description of the tests).

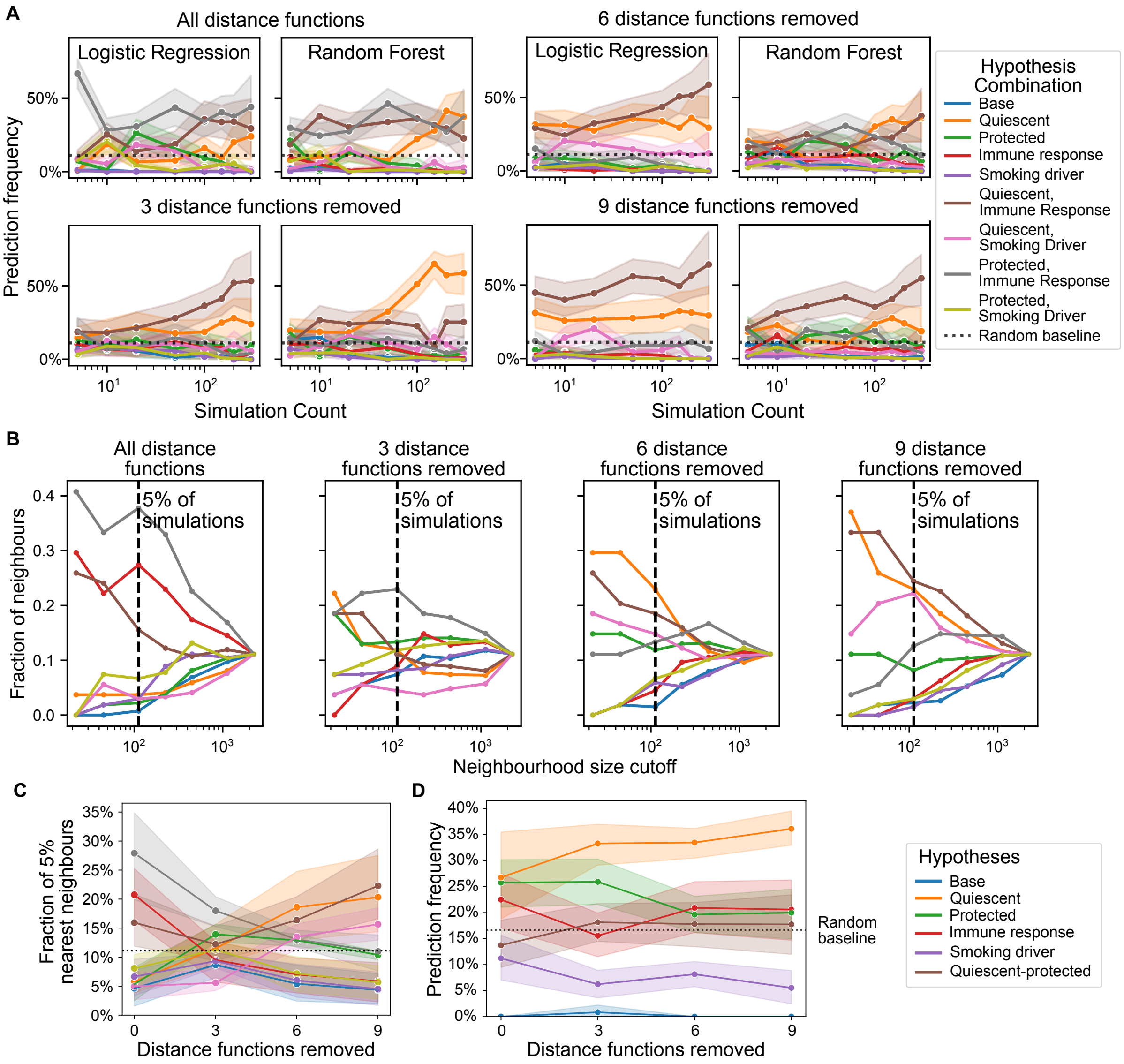


#### Figure S12. Aggregated distance classifier concurs with MDS-based classifiers on observed data classification

(A) Prediction frequency of observed data as each paradigm (colour) by MDS-based classifiers at different observed data extremity thresholds and training simulation subsample sizes. Points represent means, lines show interpolation, and shading shows a bootstrapped 95% CI over random subsamples of training simulations.

(B) Prediction frequency of observed data by aggregated distance k-NN classifiers at different observed data extremity thresholds and nearest-neighbour thresholds.

(C) As in (B), but showing only the value at 5% for each threshold for observed data extremity. The quiescent and immune response paradigm, along with the paradigm of a quiescent subpopulation alone, are most commonly classified at the most stringent threshold.

(D) Prediction frequency of observed data among all 24 hypothetical paradigms, shown by frequency of each hypothesis' inclusion. Points and lines are as in (A), while shaded areas here reflect a bootstrapped 95% CI over technical replicate classifiers. The quiescent hypothesis is consistently favoured.

| Name | Abbrev. | Brief Summary |
|---|---|---|
| Quiescent | q | A sub-population of basal cells are of a quiescent cell type, which divide slower to produce regular basal cells |
| Quiescent-protected | qp | The quiescent sub-population are less affected by the presence of tobacco smoke (this is a sub-hypothesis) |
| Protected | p | A sub-population of cells are less affected by the presence of tobacco smoke |
| Differential Immune Response | ir | The immune system preferentially kills highly mutated cells, and is dampened during smoking |
| Smoking-addicted Drivers | sd | Mutations with positive fitness (causing proliferation within the stem cell compartment via cell fate bias) are super-charged by the presence of smoke |

**Table S2. Modelling hypotheses** Reference table for modelling hypotheses, with name, abbreviated name and a brief summary of each.

| Parameter name | Varying scale | Prior mean | Prior std |
|---|---|---|---|
| non-smoking mutations per year | log | 25 | 20 |
| smoking mutation rate augmentation | log | 3 | 2 |
| non-smoking divisions per year | log | 33 | 15 |
| smoking division rate augmentation | log | 0.8 | 0.4 |
| quiescent fraction | logit | 0.1 | 0.08 |
| quiescent divisions per year | log | 4 | 3.5 |
| ambient quiescent divisions per year | log | 16.5 | 16.5 |
| quiescent protection coefficient | logit | 0.5 | 0.4 |
| protected fraction | logit | 0.25 | 0.2 |
| protection coefficient | logit | 0.5 | 0.4 |
| immune death rate | log | 0.00025 | 0.00025 |
| smoking immune coefficient | log | 0.005 | 0.004 |
| smoking driver fitness augmentation | log | 3 | 2 |

**Table S3. Parameter prior distributions** Parameters used in the simulation model, with prior distributions derived from literature (see Methods).

| Name | Description |
|---|---|
| Wasserstein | Wasserstein distance between distributions of mutational burden |
| Wasserstein with smoking | Sum of 1D 1-Wasserstein distances of smoking- and non-smoking-associated mutational burden |
| Smoking signature | 1-Wasserstein distance between distributions of smoking-associated mutational burden |
| Z-normalised | 1-Wasserstein distance between z-score normalised distributions of mutational burden |
| Mean-normalised | 1-Wasserstein distance between distributions of mutational burden, after additive normalisation |
| Mean-normalised with smoking | 2D Wasserstein simplified distance after mean-normalisation of each distribution |

**Table S4. Mutational burden-based metrics** Reference table for mutational burden-related distance functions.

| Name | Description |
|---|---|
| Bimodality | Difference between the larger weights (absolute or squared). |
| Dominant mean | Squared difference between the mean values of the components with higher weight |
| More-mutated mean | Squared difference between the mean values of the components with larger mean |
| Less-mutated mean | Squared difference between the mean values of the components with smaller mean |
| Weighted means by Dominance | Difference between components' mean values, matched by higher/lower weight, weighted by the geometric mean of the weights |
| Weighted means by Position | Difference between components' mean values, matched by higher/lower mean, weighted by the geometric mean of the weights |

**Table S5. Gaussian mixture model-based metrics** Reference table for distance functions based on one- or two-component Gaussian mixture models fit to mutational burden distributions. BIC is used to select component number.

| Name | Description |
|---|---|
| Branch lengths (Wasserstein) | 1-Wasserstein distance between the distributions of branch lengths in the two samples' phylogenetic trees, each subsampled down to the number of cells sequenced for that patient |
| Tree balance | Difference (absolute or squared) between values of the tree balance metric J1 in the two samples' phylogenetic trees, each subsampled down to the number of cells sequenced for that patient |
| Zero control | Negative control distance, always returning 0 |
| Random control | Negative control distance, returning a random number between 0 and 1 |

**Table S6. Phylogeny-based and negative control metrics** Reference table for phylogeny-related distance functions.

| VIF threshold | Selected independent distance functions |
|---|---|
| 20 | Wasserstein, Smoking signature, Z-normalised, Mean-normalised with smoking, Bimodality (squared), More-mutated mean, Less-mutated mean, Weighted mean by position, Total branch length (later omitted), Tree balance (squared) |
| 5 | Smoking signature, Z-normalised, Mean-normalised with smoking, Bimodality (squared), Less-mutated mean, Weighted mean by position, Tree balance (squared) |

**Table S7. Metrics remaining after the VIF reduction greedy search to** 20% **and** 5% "Most independent" subset of distance functions, after stopping the greedy search algorithm at maximal VIF 20 or 5.

| Parameter name | Prior mean | Posterior mean | Adjusted p-value |
|---|---|---|---|
| Non Smoking Mutations Per Year | 25 | 46.099377 | 2.493788e-156 |
| Smoking Mutation Rate Augmentation | 3 | 2.746913 | 7.999657e-03 |
| Fitness Change Scale | 0.0375 | 0.045700 | 2.389850e-18 |
| Non Smoking Divisions Per Year | 33 | 30.969207 | 7.522469e-03 |
| Smoking Division Rate Augmentation | 0.8 | 0.791367 | 9.121819e-01 |
| Quiescent Fraction | 0.1 | 0.100645 | 2.974297e-02 |
| Quiescent Divisions Per Year | 4 | 4.381729 | 2.861816e-02 |
| Ambient Quiescent Divisions Per Year | 16.5 | 13.300004 | 2.485812e-05 |
| Quiescent Gland Cell Count | 11.989 | 6.752000 | 2.389850e-18 |
| Immune Death Rate | 0.00025 | 0.000253 | 1.635272e-01 |
| Smoking Immune Coeff | 0.005 | 0.005051 | 3.574733e-01 |

**Table S8. Significance testing of parameter value selection in simulations close to observed data** Comparison of parameter values between prior distributions and 5% closest simulations to the true data. 10,000 simulations were run in the quiescent and immune response paradigm with parameter values drawn from prior distributions. Distances were calculated by normalised aggregated distance functions after the strictest filtration for observed data extremity. Kolmogorov-Smirnov tests were run for parameters with continuous prior distributions, and Chi-squared tests for parameters with discrete prior distributions. P-values were corrected for multiple testing via the Benjamini-Hochberg procedure.
